# Malignant astrocyte swelling and impaired glutamate clearance drive the expansion of injurious spreading depolarization foci

**DOI:** 10.1101/2020.10.02.324103

**Authors:** Ákos Menyhárt, Rita Frank, Attila E. Farkas, Zoltán Süle, Viktória É. Varga, Ádám Nyúl-Tóth, Anne Meiller, Orsolya Ivánkovits-Kiss, Coline L. Lemale, Írisz Szabó, Réka Tóth, Dániel Zölei-Szénási, Johannes Woitzik, Stephane Marinesco, István A. Krizbai, Ferenc Bari, Jens P. Dreier, Eszter Farkas

## Abstract

Spreading depolarizations (SD) indicate infarct maturation and predict worse clinical outcome in ischemic stroke. We demonstrate here in rodents that brain edema formation upon ischemic stroke impairs astroglial glutamate clearance and increases the tissue area invaded by SD. The cytotoxic glutamate accumulation predisposes an extensive bulk of tissue for a yet undescribed simultaneous depolarization (SiD). We confirm in rat brain slices under hypo-osmotic stress that SiD is the pathological expansion of prior SD foci, is associated with astrocyte swelling and triggers oncotic neuron death. The blockade of astrocytic aquaporin-4 channels and Na^+^/K^+^/Cl^-^ co-transporters, or volume-regulated anion channels mitigated slice edema, glutamate accumulation and SiD occurrence. Reversal of slice edema by hyperosmotic treatment counteracted glutamate accumulation and prevented SiD. In contrast, paralysis of astrocyte metabolism or inhibition of astrocyte glutamate uptake reproduced the SiD phenotype. We discuss our results in the light of evidence for SiD in the human cortex. Our results emphasize the need of preventive osmotherapy in ischemic stroke.

## Introduction

Cerebral edema is a key prognosticator of unfavorable outcome in acute ischemic, hemorrhagic or traumatic brain injury, in which spreading depolarizations (SDs) are implicated. Moreover, the robust correlate of lesion progression after acute brain injury is a characteristic pattern of SDs, which serves as an electrophysiological biomarker of injury progression, and is considered as a target of pharmacological intervention (Lauritzen et al., 2011, Dreier et al., 2017, Hartings et al., 2017; Klass et al., 2018). SD is a slowly propagating wave (2-6 mm/min) of a near complete cellular depolarization followed by the transient depression of neural activity and cytotoxic edema (Somjen 2001, Dreier 2011, Dreier et al., 2018). In addition to the cytotoxic water translocation within the nervous tissue, ischemic SD initiates cerebrospinal fluid influx and drives acute brain swelling after middle cerebral artery occlusion in mice (Mestre et al., 2020).

Although SD is primarily the profound ionic disturbance of neurons (Somjen 2001), intact astrocytic clearance mechanisms are essential for the recovery of the tissue from SD. Under stress, the notable swelling of astroglia impairs K^+^ and glutamate clearance (Kimelberg and Kettenmann, 1990), which makes neurons susceptible for increased action potential firing, epileptiform activity and SD (Chebabo et al., 1995, Roper et al., 1992), and compromises neuronal viability (Rossi et al., 2007). Moreover, astrocyte swelling in response to metabolic poisoning by fluorocitrate provoked spontaneous SD occurrence, and prolonged SD duration and neuronal injury in anesthetized rats (Largo et al., 1996, Largo et al., 1997; Larrosa et al., 2006). In focal cerebral ischemia, the degree of astrocyte soma swelling was found coincident with the cumulative duration of recurrent SDs and dendritic beading (Risher et al., 2010; Risher et al., 2012).

SD has been recognized to propagate from a punctual focus, yet the cellular mechanistic understanding of SD eruption is incomplete. The minimum tissue volume of the SD focus was estimated to be ~1 mm^3^ *in vivo*, or even smaller, ~0.03–0.06 mm^3^ *in vitro* (Matsuura and Bures, 1971, Verhaegen et al., 1992; Tang et al., 2014, Hartings et al., 2017). In focal ischemia, this critical mass of tissue is localized to the inner penumbra, where instable or metastable hot zones created by metabolic supply-demand mismatch give rise to SD (von Bornstädt et al., 2015, Hartings et al., 2017). In these particular hot zones, astrocytic K^+^ and glutamate uptake may be significantly arrested (Seidel et al., 2015), which is expected to compel a contiguous group of equally exhausted, ATP depleted neurons to fail to maintain the resting ion gradients across their cell membrane, and lose their resting potential instantaneously. This simultaneously depolarized, discrete tissue volume is suggested to correspond to the SD focus. The spatial characteristics and the pathophysiological significance of the SD focus have remained poorly understood, because the focus of a spontaneous SD has been rarely captured due to its spatiotemporal unpredictability (Bere et al., 2014; Tang et al., 2014).

Here we present an original observation in the anesthetized rat that the simultaneously depolarized tissue volume denoting the focus of an SD event (i.e. simultaneous depolarization, SiD) may become extensive rather than punctual in the severely challenged, anoxic/ischemic cerebral cortex. SiD was associated with brain edema formation, and malignant glial swelling. Further, SiD encompassed tissue that previously participated in the propagation of an SD, and aggravated histological damage. Next, glutamate accumulation linked to astrocyte swelling appeared to foster SiD occurrence. Pharmacological attenuation of glial swelling and glutamate accumulation, or hyperosmotic treatment prevented SiD. We also observed SiD in the water intoxication model of cytotoxic edema, where astrocytes were permanently swollen. Finally, our results are discussed in the light of evidence for SiD in the human cortex.

## Materials and Methods

The experimental procedures were approved by the National Food Chain Safety and Animal Health Directorate of Csongrád-Csanád County, Hungary. The procedures were performed according to the guidelines of the Scientific Committee of Animal Experimentation of the Hungarian Academy of Sciences (updated Law and Regulations on Animal Protection: 40/2013. (II. 14.) Gov. of Hungary), following the EU Directive 2010/ 63/EU on the protection of animals used for scientific purposes, and reported in compliance with the ARRIVE guidelines.

### Global forebrain ischemia/anoxia in anesthetized rats

Adult, male Sprague-Dawley rats (Charles River Laboratories, 18 months old, 645±100g, n=17; electrophysiology and imaging: 12 rats, glutamate biosensors and histology: 5 rats) were used in this study. Standard rodent chow and tap water were supplied *ad libitum*. The animals were housed under constant temperature, humidity, and lighting conditions (23 °C, 12:12 h light/dark cycle, lights on at 7 a.m.). Animals were anesthetized with 1.5-2 % isoflurane in N_2_O:O_2_ (70%:30 %), and allowed to breathe spontaneously through a head mask throughout the experiment. Body (core) temperature was kept at 37 °C with a feedback-controlled heating pad. Atropine (0.1 %, 0.1 ml) was administered intramuscularly shortly before surgical procedures to avoid the production of airway mucus. Mean arterial blood pressure (MABP) was monitored continuously, and arterial blood gases were checked regularly (i.e. during baseline, under ischemia, and under anoxia) via a catheter inserted into the left femoral artery. Level of anesthesia was controlled with the aid of MABP displayed live as experiments were in progress.

A midline incision was made on the neck and both common carotid arteries were gently separated from the surrounding tissue and the vagal nerves. Lidocaine (1%) was applied topically before opening the tissue layers during preparation. A silicone coated fishing line used as occluder was looped around the common carotid arteries for the later induction of cerebral ischemia.

For electrophysiological data acquisition and imaging, a large craniotomy (4 × 5 mm) was prepared in the right parietal bone using a dental drill (ProLab Basic, Bien Air 810, Switzerland). The dura in the craniotomy was carefully removed, and the exposed brain surface was regularly rinsed with artificial cerebrospinal fluid (aCSF; mM concentrations: 126.6 NaCl, 3 KCl, 1.5 CaCl_2_, 1.2 MgCl_2_, 24.5 NaHCO_3_, 6.7 urea, 3.7 glucose bubbled with 95 % O_2_ and 5 % CO_2_ to achieve a constant pH of 7.4). For imaging, the cranial window was closed by a microscopic cover glass to serve *in vivo* optical imaging as described earlier (Farkas et al., 2008; Obrenovitch et al., 2009).

Incomplete global forebrain ischemia was induced by the permanent occlusion of both common carotid arteries (“2-vessel occlusion”, 2VO), which readily gave rise to a single spontaneous SD. In one rat, no SD emerged in response to 2VO, therefore that particular experiment was not taken for further analysis. Oxygen was withdrawn from the anesthetic gas mixture 50 min later to create an episode of anoxia, which produced anoxic/ischemic depolarization (recurrent SD: rSD, or simultaneous depolarization: SiD). The condition of anoxia was resolved by re-oxygenation 5 min later, followed by an additional 25 min of data acquisition.

#### Electrophysiology and VS dye imaging

Local field potential (LFP) filtered in direct current (DC) potential mode, and local changes of cerebral blood flow (CBF) were acquired at two distinct locations (inter-electrode distance of 4-4.5 mm). Two saline-filled (120 mM NaCl) glass capillary microelectrodes (20 μm outer tip diameter) were inserted 700 μm deep into the right somatosensory cortex. An Ag/AgCl electrode placed under the skin of the animal’s neck was used as common ground. The microelectrode was connected to custom-made dual-channel electrometer (including AD549LH, Analog Devices, Norwood, MA, USA) via Ag/AgCl leads. The recorded LFP signal was further amplified and filtered by dedicated differential amplifiers and associated filter modules (NL106 and NL125, NeuroLog System, Digitimer Ltd., United Kingdom). The recorded analogue signal was converted and displayed live using a dedicated analog-to-digital converter (MP 150, Biopac Systems, Inc.) at a sampling frequency of 1 kHz. Local CBF changes were monitored with two laser-Doppler needle probes (Probe 403 connected to PeriFlux 5000; Perimed AB, Sweden) positioned next to the penetration site of the LFP microelectrodes. The LDF signal was digitalized and acquired, together with the DC potential, essentially as described above.

A voltage-sensitive dye (VS dye; RH-1838, Optical Imaging Ltd, Rehovot, Israel) was dissolved in aCSF, and circulated in the closed cranial window to saturate the cortical tissue as reported earlier (Farkas et al., 2008). For live imaging, the cortex was illuminated in stroboscopic mode (100 ms/s) with a high-power light emitting diode (LED) (625 nm peak wavelength; SLS-0307-A, Mightex, Pleasanton, CA, USA) equipped with an emission filter 620–640 nm bandpass; 3RD620 640, Omega Optical Inc. Brattleboro, VT), and with a laser diode (HL6545MG, Thorlabs Inc., New Jersey, USA; 120 mW; 660 nm emission wavelength) driven by a power supply (LDTC0520, Wavelength Electronics, Inc., Bozeman, USA) set to deliver a 160-mA current (Obrenovitch et al., 2009; Farkas et al., 2010; Menyhárt et al., 2017). Two identical CCD cameras (resolution: 1024 × 1024 pixel, Pantera 1M30, DALSA, Gröbenzell, Germany) were attached to a stereomicroscope (MZ12.5, Leica Microsystems, Wetzlar, Germany) equipped with a 1:1 binocular/video-tube beam splitter. VS-dye fluorescence was captured with a camera equipped with a bandpass filter (3RD 670–740; Omega Optical Inc.). In order to create CBF maps by laser speckle contrast analysis (LASCA), raw speckle images were captured by the second CCD camera (1 frame/second; 2 ms for illumination and 100 ms for exposure). A dedicated program written in LabVIEW environment synchronized the illumination and camera exposures. CBF maps were generated from the corresponding raw images as reported earlier (Menyhárt et al., 2017). Changes in VS dye fluorescence intensity and CBF with time were extracted by placing regions of interest (ROIs) of 19 × 19 pixel size (~70 × 70 μm) at selected sites on the surface of cerebral cortex.

#### Measurement of extra-synaptic glutamate concentration

Extra-synaptic glutamate concentrations were acquired using oxidase enzyme-based microelectrode biosensors (tip diameter: 30-40 μm) with constant potential amperometry (Vasylieva et al., 2013). Biosensors were constructed of Pt/Ir wires (Goodfellow, Huntington, UK), inserted into a pulled glass capillary (Harvard Apparatus, Edenbridge, UK). A poly-m-phenylenediamine (PPD) screening layer was deposited to the Pt/Ir surface by electropolymerization (20 mns) with 700 mV constant potential. An encapsulated bio-layer containing glutamate oxidase was then deposited and fixed on the Pt/Ir surface. Biosensors were calibrated before and after animal or slice experiments in 0.01 M phosphate-buffered saline (PBS) with stepwise injections of glutamate concentration standards (5, 10, 15, 20, 25, 30, 35, 40, 45 μM). Glutamate oxidase enzyme sensitivity was confirmed by the application of 5 μM D-serine. During measurements, a constant potential of 500 mV was applied vs. an Ag/AgCl-electrode placed in the recording tissue chamber or under the skin of the animal’s neck. Glutamate biosensors were lowered into the cortex together with a control sensor covered with bovine serum albumin only (BSA, SigmaAldrich, St Quentin Fallavier, France). These control biosensors recorded negligible currents (glutamate independent) compared with glutamate biosensors. Biosensors were connected to a dedicated 3-electrode potentiostat (Quadstat) equipped with an eDAQ data acquisition system (eDAQ Pty Ltd., Colorado Springs, CO, USA) and a dedicated eDAQ Chart program.

#### Histology

Animals were transcardially perfused with physiological saline followed by 4 % paraformaldehyde (PFA). The brains were removed and post-fixed in 4 % PFA for 24 hours. 20 μm brain slices were cut with a freezing microtome after cryoprotection with 30 % sucrose in PBS.

Animals were transcardially perfused with physiological saline followed by 4 % paraformaldehyde (PFA). The brains were removed and post-fixed in 4 % PFA for 24 hours. 20 μm brain slices were cut with a freezing microtome after cryoprotection with 30 % sucrose in PBS.

Nissl staining to visualize necrosis in the cortex of anesthetized rats was carried out on 20 μm brain slices mounted on Superfrost+ slides in 0.3 % polyvinyl-alcohol solution and dried at room temperature. Slides were incubated with 1 % cresyl violet (containing 0.08 % acetic acid) for 20 min at 50°C, rinsed with distilled water, dehydrated, and mounted with Depex^®^ mounting medium. The slides were examined under a Nikon Eclipse 80i brightfield microscope at 4 x and 20 x magnification, and images were captured with a camera (QImaging MicroPublisher) operated via ImagePro Plus software.

For fluorescent immunocytochemical labeling, antigens were retrieved with citrate buffer for 10 min at 95°C. The slices were permeabilized with 0.2 % TritonX-100 in PBS for 30 min, then blocked with 3 % BSA in PBS for 1 h at room temperature and incubated with primary antibodies (mouse anti-GFAP, Sigma, 1:500; mouse anti-NeuN, Millipore, 1:500; rabbit anti-cleaved caspase-3, Cell Signaling, 1:500) overnight in 3 % BSA in PBS at 4°C. The slices were incubated with secondary antibodies (goat anti-mouse Alexa-488, Jackson Laboratories, 1:500; goat anti-rabbit Cy3, Jackson Laboratories, 1:500; goat anti-mouse Star Red, Abberior, 1:500 for STED) in 1 % BSA in PBS for 1 h at room temperature. Finally, cell nuclei were labelled with Hoechst staining (1:1000) for 10 min. The slices were mounted with Fluoromount-G and stored at 4°C. Photomicrographs were taken (3 per animal per each selected brain region of the hemisphere ipsilateral to the craniotomy) with a Nikon Eclipse TE2000U microscope at 20 x magnification. Regions of interest, expected to be most vulnerable to ischemia/anoxia were: superficial and deeper layers of cortex, hippocampal CA1 and striatum. Representative confocal images were taken with Leica SP5 inverted laser scanning confocal microscope at 20 x magnification (Z-stacks with steps of 0.4 μm). STED images were acquired with a Stedycon STED instrument (Abberior Instruments, Göttingen, Germany) attached to a Zeiss Axio Observer Z1 inverted microscope. Confocal and STED images were pieced together and filtered in ImageJ. The distinctive tissue autofluorescence of brain samples was corrected by subtracting images captured in the non-labelled far red spectrum from the respective confocal photomicrographs. NeuN and GFAP immunopositive cells, alone and co-localized with cleaved caspase-3 were counted manually with Cell Counter plugin of the Image J software by two independent observers.

### Hypo-osmotic cerebral edema model in anesthetized rats and mice

#### Water intoxication in anesthetized rats: electrophysiology

Male Wistar rats (Charles River Laboratories, 2 months old, 250±100g, n=11) were used. Anesthesia and surgical procedures (including craniotomy) were identical to those presented above, except for carotid artery preparation and occlusion. In brief, the rostral craniotomy incorporated two LFP electrodes to measure SD evolution, while the caudal craniotomy served SD elicitation. Hypo-osmotic brain edema was achieved by water intoxication imposed by intraperitoneal injection of distilled water (DW; 15 % body weight) (Yamaguchi et al., 1994, n=6). In the control group (n=4) physiological saline was administered in the same volume as DW. In one rat, SD occurred spontaneously due to DW administration. In the other rats (n=6), continuous SD elicitation was achieved by placing and leaving a 1 M KCl soaked cotton ball to the brain surface through a smaller trepanation drilled rostral to the first craniotomy, after 30 minutes of DW or physiological saline administration. KCl was washed out with aCSF after the occurrence of the first SD, and 30 min later O_2_ was withdrawn from the anesthetic gas mixture for 5 minutes. MABP was monitored continuously, and arterial blood gases were checked regularly via a catheter inserted into the left femoral artery. Level of anesthesia was controlled with the aid of MABP displayed live as experiments were in progress. Full data analysis was carried out in the experiments, in which SD1 was elicited with KCl.

#### Water intoxication in anesthetized mice: two-photon imaging

Male C57BL/6 mice (8-10 weeks old, n=10) were anesthetized with 1% Avertin (20 μL/g, i.p.), and mounted on a stereotactic frame incorporating a heating pad (Menyhárt et al., 2018). A cranial window (d=3 mm) was prepared on the right parietal bone, and the dura was retracted. For astrocyte calcium imaging, the exposed brain surface was first loaded topically with a green fluorescent calcium indicator Fluo 4-AM (45 μM in aCSF, Thermo Fisher, Waltham, MA, USA) and incubated for 15 minutes. Subsequently, to monitor astrocyte swelling, the astrocyte specific non-fixable red fluorescent dye sulforhodamine 101 (SR101, 80 μM in aCSF, Thermo Fisher, Waltham, MA, USA) was applied topically, and incubated for further 15 minutes. The craniotomy was then closed with a microscopic cover glass. Multiphoton excitation was performed at 810 nm wavelength according to the protocols described previously (Menyhárt et al., 2018; Supplement to this paper). Final imaging depth in the parietal cortex was 55-85 μm, where a z-stack with 5 μm vertical steps was recorded at the area of interest for identification of astrocytes. Image sequences were taken of the desired cells at approximately 1 μm/pixel spatial and 0.8–2.5 Hz temporal resolution. After baseline (5 min), mice received stepwise intraperitoneal DW injections (3 times 1.5 ml/10 minutes) to 15 % volume of body weight. SD was then elicited with the topical application of 1 M KCl, and SD evolution was confirmed by the occurrence of synchronous astrocytic calcium waves (Fluo 4-AM intensity increase, green channel) and the associated volume changes of astrocyte soma and processes (SR101 labeled soma volume changes, red channel).

### Hypo-osmotic cerebral edema model in live rat brain slice preparations

#### Brain slice preparation, incubation media and experimental protocols

Adult male Wistar rats (body weight: 250 g; n=47) were decapitated under deep anesthesia (4-5 % isoflurane in N_2_O:O_2_; 2:1), and coronal brain slices were prepared as reported previously (Menyhárt et al., 2018). Briefly, coronal brain slices (350 μm) anterior to bregma were cut with a vibrating blade microtome (Leica VT1000S), and collected in ice-cold aCSF (composition of aCSF in mM concentrations: 130 NaCl, 3.5 KCl, 1 NaH_2_PO_4_, 24 NaHCO_3_, 1 CaCl_2_, 3 MgSO_4_ and 10 D-glucose). Six to eight slices were transferred to an incubation chamber filled with carbogenated aCSF (difference in composition in mM concentrations: 3 CaCl_2_ and 1.5 MgSO_4_), and kept at room temperature (~20°C). Selected slices were placed into an interface type recording tissue chamber (Brain Slice Chamber BSC1, Scientific Systems Design Inc., Ontario, Canada) continuously perfused with carbogenated aCSF at a rate of 2 ml/min, and kept at 32°C using a dedicated proportional temperature controller unit (PTC03, Scientific Systems Design Inc., Ontario, Canada).

Hypo-osmotic solutions were prepared by reducing the NaCl concentration of aCSF from the regular 130 to 40-120 mM (HM_40_-HM_120_), while other components and the pH of the medium were unaltered. The results shown in this paper were obtained with HM_100_ and HM_60_. The use of HM_60_ was relevant for the pharmacological set of experiments, because this condition regularly evoked spontaneous SDs to be modified by the drugs used. Iso-osmotic hyponatremic solutions were prepared by substituting the withdrawn NaCl with equiosmolar mannitol. Hyper-osmotic solutions (HRM) contained additional mannitol in normal aCSF at 100 mM concentration (n=11). Over the experimental protocol, control slices were incubated in aCSF throughout the recording (n=21), whereas aCSF was replaced with HM (i.e. typically HM_100_ or HM_60_) in the recording chamber as the experimental condition (n=57, HM_100_: 16 slices, HM_60_: 41 slices).

In control slices incubated in aCSF, the first SD (SD1) was triggered by electric stimulation as reported earlier (Hertelendy et al., 2017). A subsequent, recurrent SD (rSD) was elicited by transient anoxia (i.e. the replacement of O_2_ with N_2_ in the gas mixture applied to bubble the aCSF for 2.5 min). Under HM, SD1 typically occurred spontaneously in response to osmotic stress, and the subsequent event was elicited 15 min later with transient anoxia (Fig. 2).

**Figure 1.**
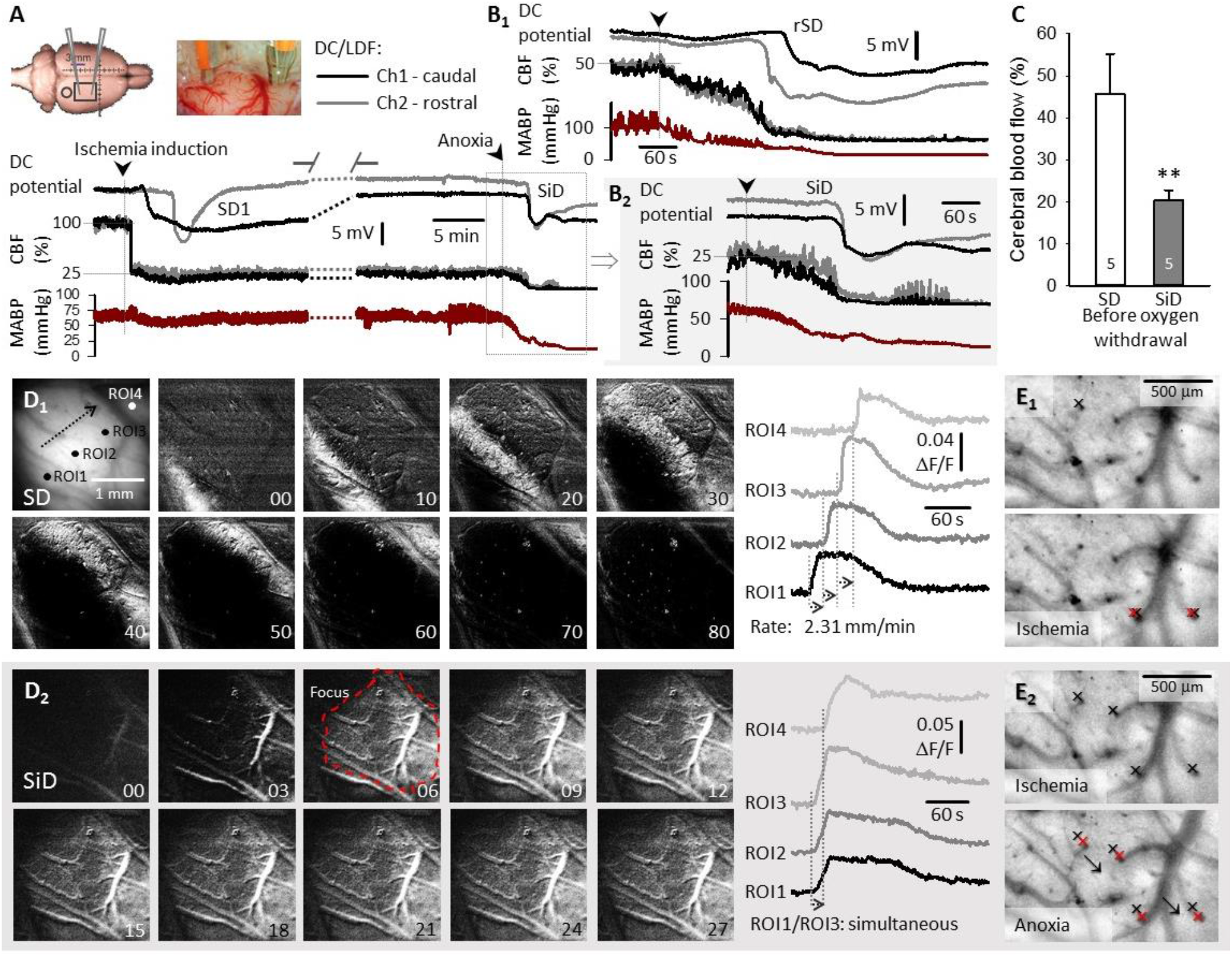
The occurrence of SD and SiD with respect to the degree of hypoperfusion and tissue swelling in the cerebral cortex of anesthetized rats. **A,** A representative recording of a spontaneous SD in response to ischemia onset (SD1), and a subsequent SiD in response to anoxia. **B,** In 6 of 12 rats, anoxia triggered a spreading depolarization event (rSD) (B1). In the remaining 6 rats, anoxia provoked SiD (B2). **C,** CBF levels before anoxia initiation. **D,** Background subtracted voltage sensitive (VS) dye image sequences confirm SD1 propagation from the caudo-lateral towards the fronto-medial corner of the field of view after ischemia onset (D1). The arrow in the first image and traces derived from the ROIs show the propagation of SD1. In contrast, anoxia triggered an SiD that appeared at ROI1 and ROI2 in synchrony, and involved a considerable part of field of view simultaneously (D2). **E,** Tissue edema is shown with the shift of contrast edge markers on contrasted VS dye images. Note the position of the red markers after ischemia (E1) and anoxia (E2) with respect to the black markers placed prior to the challenges.

**Figure 2.**
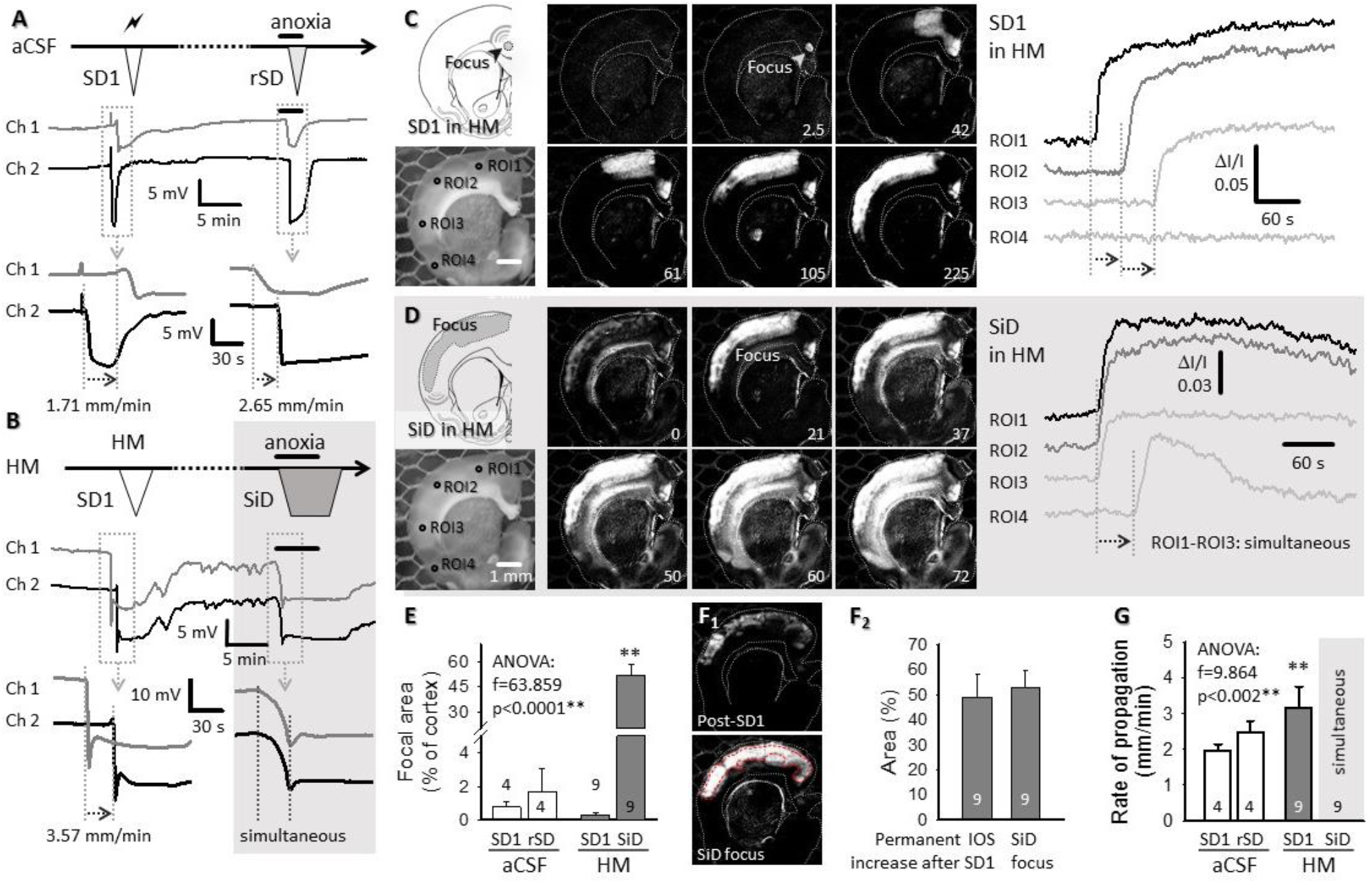
Replication of SD and SiD during osmotic stress in rat live coronal brain slice preparations. **A,** The regular spreading nature of both SD1 and rSD in normal aCSF is shown in a representative electrophysiology experiment. SD1 was elicited by electrical stimulation and rSD was induced by transient anoxia (oxygen withdrawal of 2.5. min). **B,** SD1 occurred spontaneously to HM exposure (HM60 here), while the subsequent event was induced 15 min later by transient anoxia. The original electrophysiological recordings demonstrate that SD1 was a spreading event, followed by an SiD in response to anoxia. **C and D,** In background subtracted IOS images of a HM incubated brain slice, the temporal characteristics of SD1 in Panel C reveal a punctual focus and propagation in the upper cortical layers. In Panel D, subsequent anoxia in HM (HM_100_ here) induced SiD of an extensive bulk of cortical tissue (ROI1-ROI3; SiD), and the propagation of the event towards the ventral tips of the cortex (ROI3→ROI4). Note the simultaneous increase of signal intensity (Panel D) at ROI1-ROI3, and the delay at ROI4 with respect to ROI1. **E,** The relative size of the focal area of SD/SiD measured in IOS images. **F,** The focal area of SiD incorporated the tissue zone characterized by sustained IOS intensity elevation following SD1. Images in F_1_ were taken prior to SiD (post-SD1), and at the emergence of SiD (SiD focus). A red broken line delineates the post-SD1 high IOS intensity zone (SiD focus). The mean size of the tissue areas is given in F_2_ relative to the full size of the cortex. **G,** The rate of propagation of depolarization events. In Panels E-G, data are given as mean±stdev, the number of events analyzed is shown in each bar. A one-way ANOVA paradigm was used, followed by Sidak post hoc test whenever relevant (p<0.05*p<0.01**).

To characterize the excitability of the nervous tissue, evoked field potentials (EFP) were recorded from layer 3-4 of the somatosensory cortex, or CA1 region of the hippocampus. Slices were incubated in aCSF or in HM. Stimulation was delivered by a concentric bipolar needle electrode. The distance between the current delivery electrode and recording microelectrode was approximately 800 μm. Stimulation was implemented with single constant current pulses (5-15 μA, 1 ms).

Data were collected by LFP recording, optical imaging or extra-synaptic glutamate concentration measurement with enzymatic biosensors, each alone or in combination, as described below. Slices, in which spontaneous SD1 did not occur, were not taken for quantitative analysis.

#### LFP recordings

LFP measurements filtered in DC mode (<1Hz) were acquired via two glass capillary microelectrodes (1-3 MΩ) filled with 150 mM NaCl and 1 mM HEPES, and positioned at a distance of 1000 μm with respect to each other, inserted into the 3^rd^ cortical layer. An Ag/AgCl electrode was placed in the recording chamber and served as reference. Microelectrodes were connected to a custom-made dual-channel electrometer (including AD549LH, Analog Devices, Norwood, MA, USA) and dedicated differential amplifiers and associated filter modules (NL106 and NL125, NeuroLog System, Digitimer Ltd., United Kingdom). The recorded analogue signal was converted and displayed live using an Acknowledge environment (MP 150, Biopac Systems, Inc.) at a sampling frequency of 1 kHz (Menyhárt et al., 2018). The DC potential traces confirmed the occurrence of depolarization events. Further, the DC potential recordings were used off-line to characterize the spreading or simultaneous nature of depolarization events, and to determine event latency and duration at half amplitude.

EFP were recorded as described above. LFP was filtered between 1-200 Hz, and the recorded analogue signal was converted and displayed live using an Acknowledge environment. Recordings were used off-line to determine the amplitude of the evoked potentials.

#### Intrinsic optical signal imaging

For intrinsic optical signal (IOS) imaging, slices were illuminated by a halogen lamp (Volpi AG, Intralux 5100, Schlieren, Switzerland). Image sequences were captured at 1 Hz with a monochrome CCD camera (spatial resolution: 1024 × 1024 pixel, Pantera 1M30, DALSA, Gröbenzell, Germany) attached to a stereomicroscope (MZ12.5, Leica Microsystems, Wetzlar, Germany), yielding 6-10 x magnification. Changes in IOS intensity were extracted at three ROIs positioned along the propagation of SD1, and expressed relative to baseline intensity (ΔI/I). Image sequences were used off line to measure the degree of slice swelling associated with osmotic stress or depolarization events.

#### Measurement of extra-synaptic glutamate concentration in vitro

Glutamate biosensors were used and calibrated as described above for anesthetized rats. Biosensors were implanted adjacent to the LFP microelectrodes together with glutamate null or bovine serum albumin sensors (tip diameter: 30-40 μm, BSA, SigmaAldrich, St Quentin Fallavier, France). These control biosensors, coated with albumin, recorded negligible currents (glutamate independent) compared with glutamate biosensors. Biosensors were connected to a dedicated 3-electrode potentiostat (Quadstat) equipped with an eDAQ data acquisition system (eDAQ Pty Ltd., Colorado Springs, CO, USA) and a dedicated eDAQ Chart program. The magnitude of changes in extra-synaptic glutamate concentrations with depolarization events were expressed as area under the curve (AUC; μM × s).

#### Pharmacological treatments

Slices were randomly exposed to various pharmacological treatments: (i) For edema reduction, Na^+^/K^+^/Cl^-^ cotransporter blocker Bumetanide (Bum, Sigma-Aldrich; 1 mM) and the aquaporin-4 channel inhibitor TGN-020 (Tocris; 100 μM) were co-applied (n=16); (ii) To inhibit swelling related glutamate efflux via volume-regulated anion channels (VRAC), slices were exposed to the channel blocker DCPIB (Tocris; 20 μM) (n=21); (iii) N-methyl D-aspartate (NMDA) receptors were blocked by the non-competitive NMDA receptor antagonist MK-801 (Tocris; 100 μM), co-applied with the competitive α-amino-3-hydroxy-5-methyl-4-isoxazolepropionic acid AMPA/kainate receptor antagonist CNQX (Tocris; 20 μM) (n=13); (iv) TFB-TBOA (Tocris; 10 μM and 100 μM) an excitatory amino acid transporter inhibitor was washed on the slices to explore whether glutamate uptake was functional (n=4); (v) Finally, fluorocitrate (Sigma; 0.5-1 mM), a drug that disrupts the citrate cycle in astrocytes (Swanson and Graham, 1994) was applied to paralyze astrocytes (n=7). In case no spontaneous SD1 occurred, the slices were not considered for further analysis.

To evaluate the impact of HRM on the assessed variables, slices were incubated in HM first (i.e. typically HM_100_ or HM_60_) for 30 min, then HM was replaced with HRM. The first SD (SD1) occurred spontaneously in HM. A subsequent, recurrent SD (rSD) was elicited by transient anoxia (i.e. the replacement of O_2_ with N_2_ in the gas mixture applied to bubble the aCSF for 2.5 min) in HRM.

#### Histology

The size of the ischemic lesion in live brain slice preparations was determined by triphenyltetrazolium chloride (TTC) staining. The brain slices (n=12) were incubated in a 2 % solution of TTC in 0.1 M PBS for 20 minutes at room temperature. The sections were subsequently immersed and stored in 4 % PFA for 24 h. The stained sections were then mounted on microscope slides, coverslipped with glycerol, and the number of particles per 1000 μm^2^ was calculated after calibration and manual thresholding by using the automated inbuilt function “analyze particles” in FIJI.

To examine the morphology and swelling of astrocytes, a modified Golgi-Cox staining was used (Gull et al., 2015). Some brain slices were stained with FD Rapid GolgiStainTM Kit (FD Neurotechnologies, Inc., USA). The slices were postfixed in 4 % PFA containing 8 % glutaraldehyde for 24 h at room temperature. The sections were incubated in Golgi impregnation solution (equal volumes of solution A and solution B). Following impregnation, the brain slices were transferred into solution C and incubated at 4 °C for 24 h in dark. After the incubation in solution C, stained slices were dehydrated in solutions of increasing ethanol concentration (50 %, 70 % and 95 % and absolute ethanol), then mounted on microscope slides in 0.2 % polyvinyl-alcohol, and coverslipped with glycerol. Representative images were taken from the cortical layer 2-3 with Nikon-DS Fi3 camera attached to a Leica DM 2000 Led light microscope (Leica Microsystems GmbH, Germany).

Cross cortical blocks were dissected from representative brain slices for electron microscopic examination. The samples were postfixed in 3 % glutaraldehyde and 2.25 % dextran in 0.1 M phosphate buffer (pH=7.4) for a week. Semi-thin sections were cut plane on an ultramicrotome (Ultracut E, Reichert-Jung) and stained on object glasses with toluidine blue. Ultrathin sections were cut from the same blocks and collected on copper slot grids. The preparations were contrasted with 5 % uranyl acetate and Reynolds lead citrate solution. The samples were analyzed with a Jeol JEM-1400 Plus transmission electron microscope, and photographs were taken with a charge-coupled device camera.

### Data analysis

LFP and extracellular glutamate concentration for both *in vivo* and *in vitro* electrophysiology were acquired, displayed live, and stored using a personal computer equipped with dedicated software (AcqKnowledge 4.2 for MP 150, Biopac Systems, Inc., USA for LFP; and eDAQ Chart, eDAQ Pty Ltd., Colorado Springs, CO, USA for glutamate concentration). Other physiological variables (CBF, MABP) of anesthetized rodents were recorded simultaneously. Data analysis was conducted offline and was assisted by the inbuilt tools of AcqKnowledge 4.2 software. Local CBF changes along the experimental protocol were calculated based on 100 % baseline taken shortly before ischemia induction, and the residual LDF signal after anesthetic overdose, considered as biological zero. CBF changes were expressed relative to baseline (%). For both VS dye (anesthetized rats) and IOS imaging (live brain slice preparations), depolarization events were visualized after background subtraction, contrast enhancement and smoothing (3 point moving average) performed in Fiji. Traces of VS dye fluorescence or IOS intensity changes were extracted from raw image sequences at 4 ROIs positioned in the images offline, along the propagation of the first SD. Optical signal intensity changes were expressed as ΔF/F or ΔI/I, respectively. The VS dye traces were corrected for linear bleaching using exponential fitting in Fiji.

The velocity of SD propagation in the LFP experiments was calculated by taking the distance between the two LFP electrodes (typically 4-5 mm *in vivo* and 1000-1500 μm *in vitro*), and the time between the occurrence of the signature of an SD at the two electrodes. In the imaging experiments, the calculation of the velocity was aided by the visual signal, and was more accurate due to the perceptible direction of SD propagation (Fig 1–2).

In the IOS image sequences, several spatial features of SD/SiD events were measured after background subtraction and inbuilt automatic or manual thresholding in Fiji, and expressed relative to the full surface area of the cortex in the brain slice (Fig. 2–4). First, the focal area of each event was estimated (e.g. Fig. 2E). Next, the area of sustained IOS intensity elevation following SD1 was expressed (Fig. 2F_2_). Finally, the maximal cortical area covered by SD1 propagation or SiD were measured (Fig. 4A_2_).

**Figure 3.**
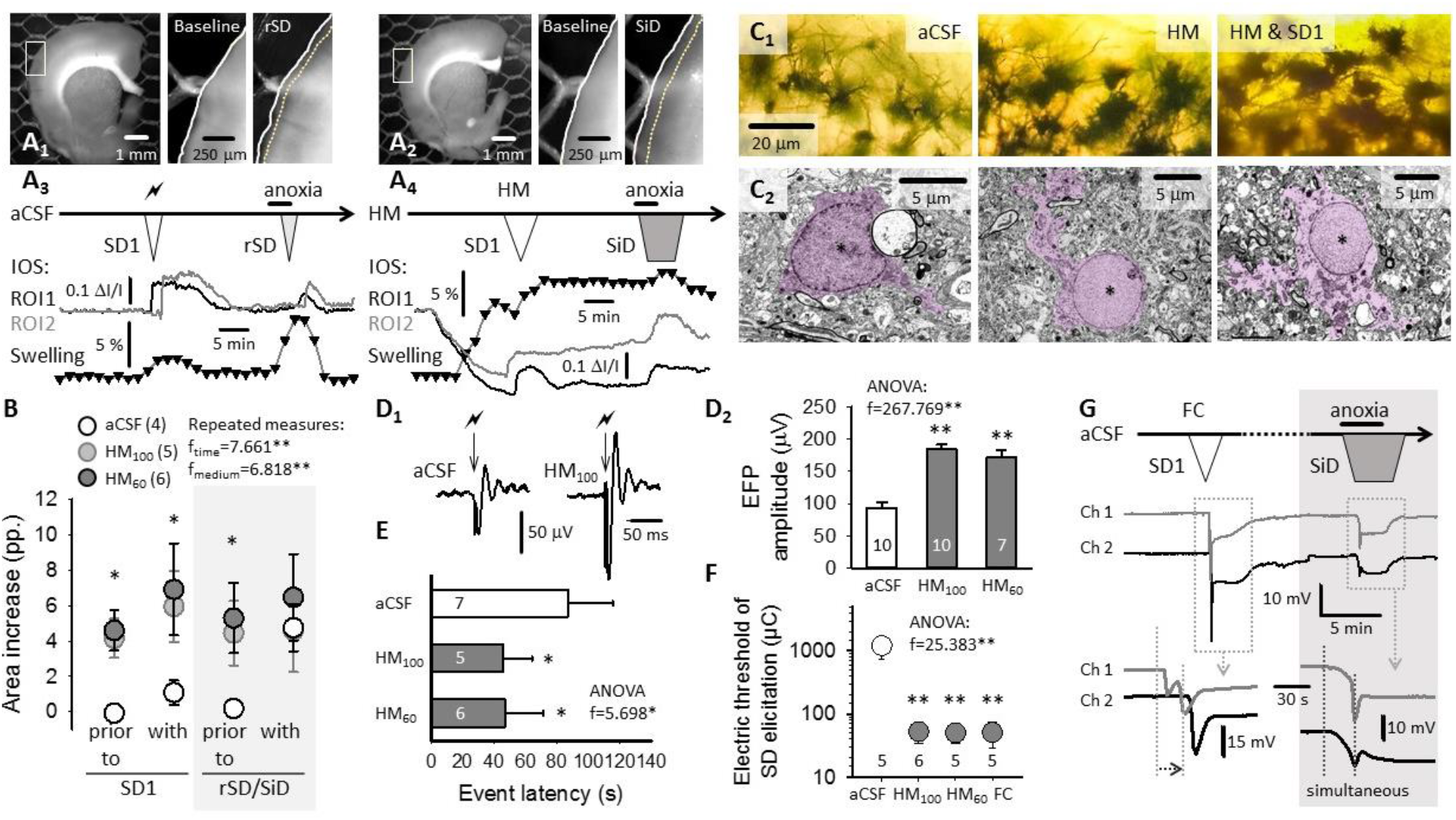
Implication of tissue edema and astrocyte swelling in SiD in brain slices. **A,** IOS images demonstrate slice swelling with rSD in normal aCSF (A_1_), and with SiD in HM (A_2_), represented by the displacement of the contour of the parieto-temporal cortex (inserts). Swelling over the experimental protocol was characterized by the change in the surface area of the full slice relative to baseline, at a temporal resolution of 1/100 s (trace with black triangles), and depicted in temporal correspondence with IOS variations at two ROIs (aCSF: A_3_; HM: A_4_). **B,** The degree of slice swelling relative to baseline at selected time points. **C,** Golgi-Cox-stained sections (C_1_) and electron photomicrographs (C_2_) of astrocytes in aCSF, HM and after SD1 in HM (asterisk: astrocyte nucleus; purple shading astrocyte plasma and nucleus). **D,** Cortical EFP amplitudes in aCSF, HM_100_ or HM_60_ (representative traces: D_1_, quantitation: D_2_). **E,** Latency of rSD in aCSF and SiD in HM after anoxia. **F,** The electric threshold of SD elicitation under aCSF, HM or fluorocitrate (FC) incubation. **G,** The application of FC to brain slices replicated the SiD phenotype observed under HM incubation. In Panels B,D_2_,-F, data are given as mean±stdev, the number of events analyzed is shown in the legend (B), each bar (D_2_ & E) or above the x-axis (F). Statistical evaluation relied on a repeated measures (B), or a one-way ANOVA paradigm (D_2_-F) (p<0.05*p<0.01**), followed by a Sidak post hoc test (p<0.05*& p<0.01** vs. aCSF).

**Figure 4.**
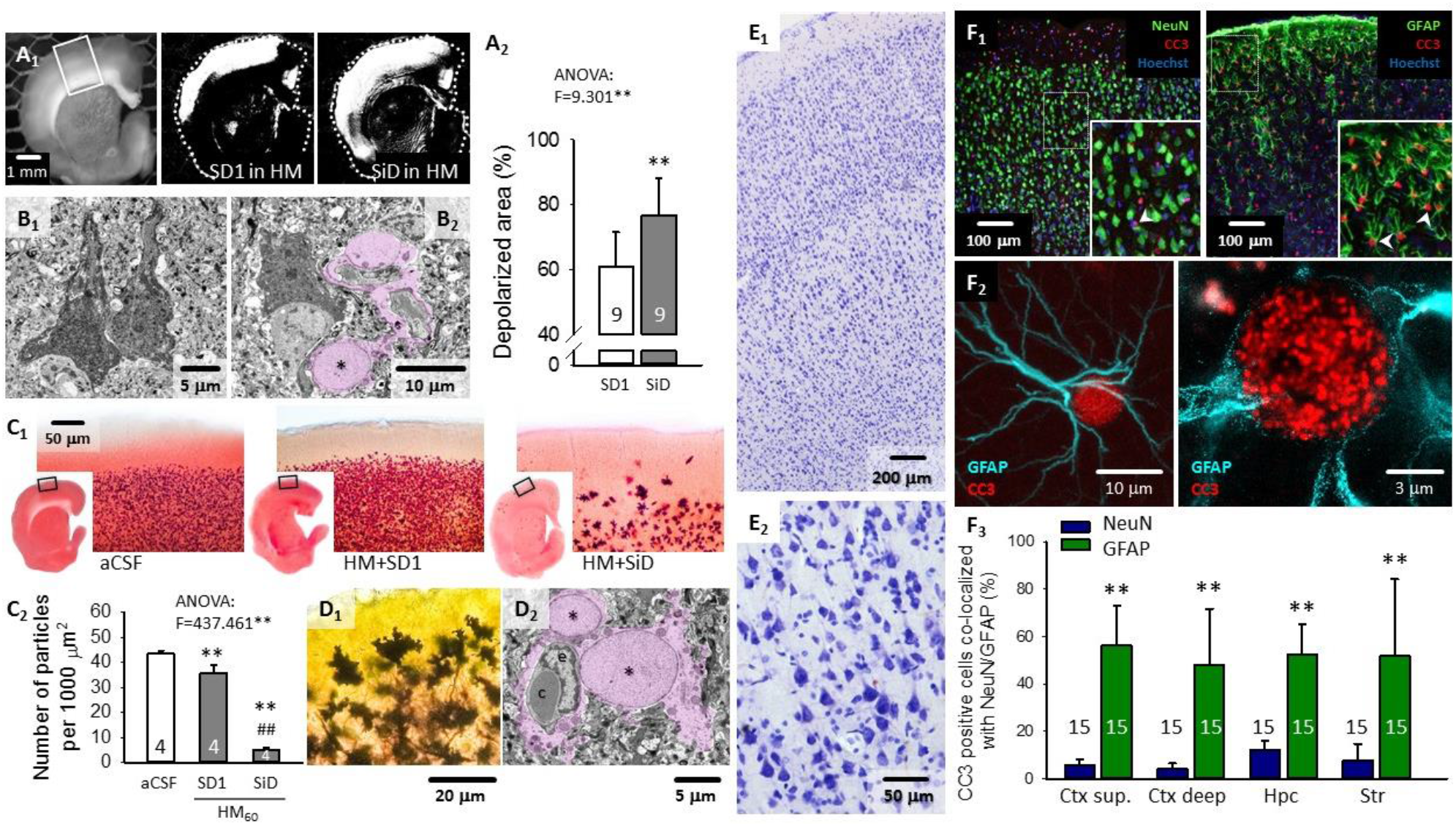
Cellular injury as a consequence of SD or SiD. **A,** The area covered by SD1 and the subsequent SiD (representative background subtracted IOS images: A_1_; quantitation: A_2_. **B,** Representative electron microscopic images show necrotic pyramidal neuronal cell death in layer 3 of the cortex (B1) and its occasional proximity to swollen astrocytes (B_2_) in brain slices after SiD in HM (asterisk: astrocyte nucleus; purple shading: astrocyte plasma and nucleus). **C,** TTC staining of brain slices after SD1 in aCSF, HM, and SiD in HM (C_1_). The number of TTC-stained cellular compartments (i.e. particles) after depolarization events (C_2_). **D,** Golgi-Cox-staining (D_1_) and electron photomicrographs (D_2_) of swollen astrocytes (asterisk, astrocyte nucleus; e, endothelial cell nucleus; c, capillary; purple shading: astrocyte plasma and nucleus). **E,** Pyramidal cell necrosis after SiD, visualized in the parietal cortex of anesthetized rats with Nissl staining. **F,** Immunocytochemical co-localization of cleaved caspase-3 (CC3) with cell nuclei (Hoechst) of neurons (NeuN) and astrocytes (GFAP) shows predominant glial apoptosis after SiD (F_1_). Super-resolution (STED) microscopy unravels the nuclear localization of CC3 and the loss of GFAP in the CC3-positive soma. Quantitative analysis of astrocyte apoptosis in the cerebral cortex (Ctx), hippocampus (Hpc) and striatum (Str) (F_3_). Data in A_2_, C_2_ and F_3_ are given as mean±stdev. Statistical analysis relied on a one-way ANOVA, p<0.01**, followed by a Sidak post hoc test (A_2_: p<0.01** vs. SD1; C_2_: p<0.01** vs. aCSF, p<0.01** vs. SD1 in HM; F_2_: p<0.01** vs. NeuN).

Brain slice swelling was shown in representative, contrasted IOS images as the 2D displacement of the contour of the parieto-temporal cortex (Fig. 3A). Slice swelling over the experimental protocol was quantitatively evaluated by the change in the surface area of the full slice relative to baseline, at a temporal resolution of 1/100 s, and depicted in temporal correspondence with IOS variations at two ROIs (Fig. 3–5). Slice area measurements were performed after contrast enhancement in Fiji.

**Figure 5.**
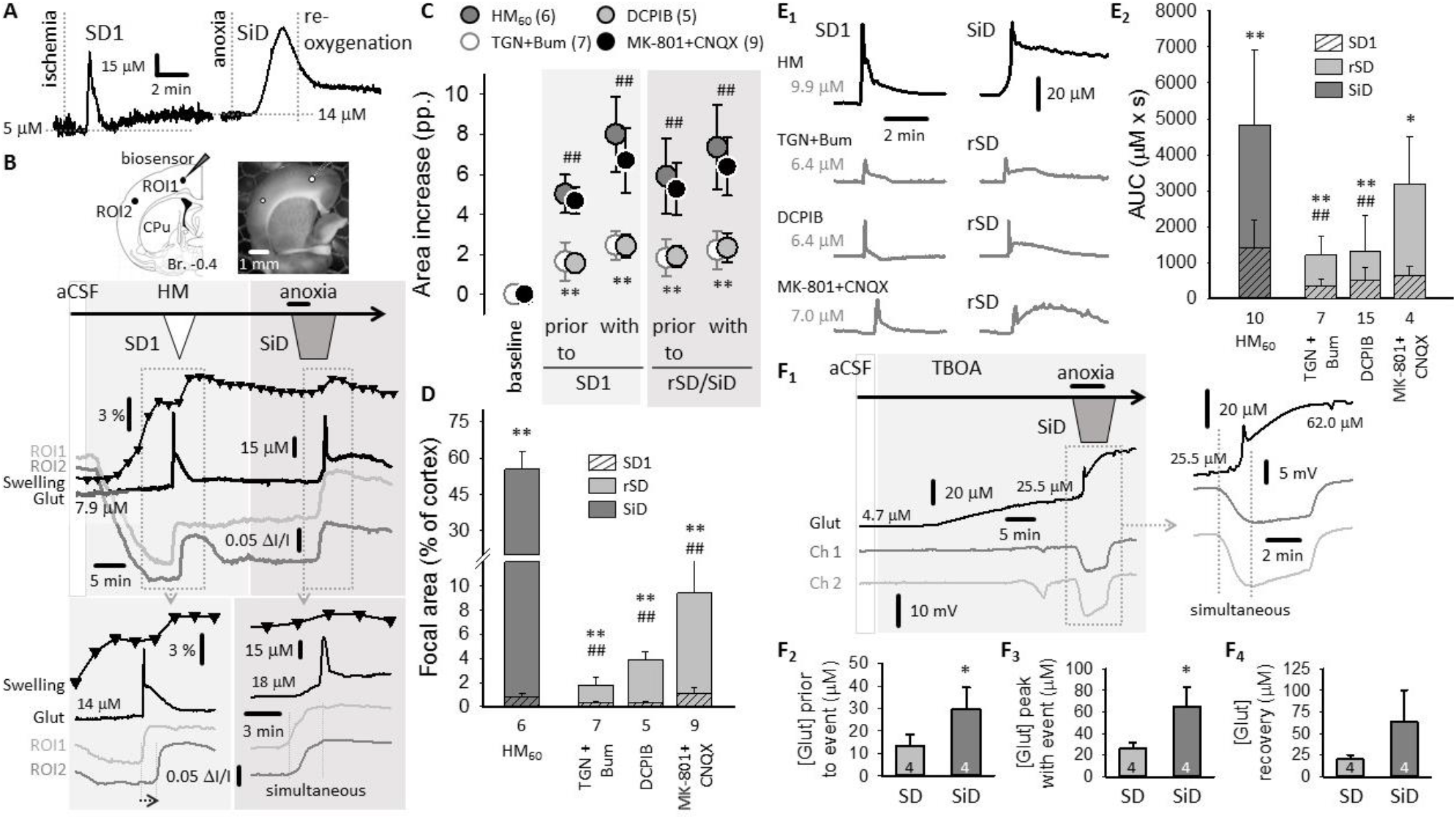
Inhibition of glial swelling or volume regulated glutamate release attenuates SiD. **A,** Transient glutamate peak with SD1 and sustained glutamate accumulation with SiD upon anoxia in an anesthetized rat. **B,** Spatiotemporal relationship between tissue swelling (black trace with black triangles), depolarization events (IOS intensity changes at ROI1 and ROI2, gray traces), and changes in glutamate concentration (black trace) in brain slices. **C,** The impact of pharmacological treatments on slice swelling relative to baseline at selected time points. **D,** The impact of pharmacological treatments on the size of the focal area of SD1 and rSD/SiD relative to the area of the cortex. **E,** Representative glutamate concentration traces (E1) and quantitative analysis (E2) of glutamate accumulation with SD1 and rSD/SiD (area under the curve, AUC). **F,** Representative recordings of glutamate concentration (Glut) and DC potential (Ch1 and Ch2) under TBOA treatment – note the occurrence of SiD (F_1_), and glutamate concentration at three selected phases of these experiments (F_2-4_). Data in C, D, E_2_ and F_2-4_ are given as mean±stdev. Statistical analysis was conducted with a repeated measures (C: Ftime=196.228**; Ftreat=29.805**) or two-way ANOVA (D: Fevent= 17.807**, Ftreat=12.801**; E: Fevent=531.829** Ftreat=14.074**), followed by a Sidak post hoc test (C: p<0.01** TGN or DCPIB vs. HM_60_; p<0.01## MK-801 vs. TGN+ Bum or DCPIB; D-E: p<0.05* and p<0.01** vs. SD1, p<0.05# and p<0.01## vs. HM), or one-way ANOVA (F: p<0.05*).

The excitability of the nervous tissue was characterized with the amplitude of evoked field potentials, and the electric threshold of SD elicitation. For the latter, the charge delivered was calculated as *Q*[μC]=*I*[mA] × *t*[ms], and it was raised stepwise with an interstimulus interval of 2 min until SD was observed. The threshold of SD elicitation was illustrated on a logarithmic scale (Fig. 3D-F).

Multiphoton image stacks were processed offline. Image stacks were auto leveled, background subtracted and converted to RGB color in FIJI. Images of individual astrocytes were cropped and cellular movement artefacts were corrected using the “Template Matching” plugin in FIJI. The SD associated intracellular calcium waves were measured on green fluorescent images (ΔF/F) by placing a 15×15 μm ROI on the soma of selected astrocytes. Astrocyte swelling was determined on red fluorescent image sequences by the morphometric analysis of somatic area changes in the lateral dimensions. After binary conversion, images were manually thresholded, astrocyte somatic areas were outlined, the outlined area was measured automatically using the inbuilt “analyze particles” function of FIJI and expressed as % area of baseline.

Quantitative data are given as mean±standard deviation (stdev). Statistical analysis was conducted with the software SPSS (IBM SPSS Statistics for Windows, Version 22.0, IBM Corp.). Data sets were evaluated by an independent samples T-test, a one-way analysis of variance (ANOVA), a two-way ANOVA or repeated measures, followed by a Sidak post hoc test when appropriate. Levels of significance were set at p<0.05* or p<0.01**. Distinct statistical methods are provided in each Figure legend in detail.

## Results

### Simultaneous depolarization occurs in response to anoxia in experimental global forebrain ischemia

The initiation of ischemia typical of penumbra tissue was confirmed by a sudden drop of CBF (to a minimum of 23.6±5.9 % of baseline), which was shortly followed by the DC potential signature of an SD (SD1). The event propagated over the cortex between the two electrodes (rate: 2.91±1.18 mm/min) (Fig. 1A). Fifty minutes later, upon O_2_ withdrawal, CBF decreased further from 38.8±15.8 to 18.4±6.3 %, MABP dropped from 105±2 to 67±1 mmHg, and arterial pO_2_ fell from 105±2 to 35±1 mmHg. The anoxic condition gave rise to a second depolarization event within 2 min (Fig. 1A-B). In half of the animals (n=5), the depolarization emerged virtually simultaneously at the two electrodes positioned 4-5 mm apart (Fig. 1A), which we designated as “simultaneous depolarization” (SiD) (Fig. 1A). In other rats (n=5), the depolarization to anoxia spread (recurrent SD, rSD), but traveled across the cortex twice as rapidly as SD1 (5.66±1.23 mm/min; Fig. 1B_1_). The occurrence of SiD, rather than rSD was predicted by the low local level of CBF measured 1 min prior to anoxia induction (20.2±2.5 vs. 45.6±9.6 %; for SiD vs. rSD) (Fig. 1C). To appreciate the spatial resolution of the electrophysiological signal, we reproduced the protocol with VS dye imaging. The optical imaging experiment confirmed SD1 propagation across the field of view after ischemia induction (2.31 mm/min), SD1 arriving in the cranial window from a focus out of the field of view (Fig. 1D_1_). Three of the four ROIs placed along the propagation of SD1 revealed the occurrence of SiD upon anoxia (Fig. 1D_2_; Suppl. Video 1). This bulk of tissue was perceived as an extensive SD focus, from which the depolarization spread to the fourth ROI (rate: 4.5 mm/min).

During image processing, we incidentally observed a large movement of the cortex upon anoxia, shortly before SiD occurrence, which corresponded to edema formation (Suppl. Video 2). The cortical edema, revealed by the shift of contrast edge markers in an image sequence, was typical before SiD, but was not seen prior to SD1 (Fig. 1E_1-2_). The association of cortical edema formation with SiD prompted the hypothesis that tissue swelling may be the underlying condition for SiD, which essentially represents a large SD focus, in contrast with a punctual SD origin. To explore this concept, we moved on to experiment on live brain slice preparations, which offered the controlled induction of tissue edema and the reliable visualization of SD focus.

### Hypo-osmotic stress favors the evolution of SiD

To test the hypothesis that brain edema formation expands the SD focus to be detected as SiD, we created hypo-osmotic stress in live brain slices, and monitored SD with two DC potential electrodes (about 1.5 mm apart) or IOS imaging. While bipolar electric stimulation was required for SD1 induction in aCSF (i.e. control slices), SD1 occurred spontaneously 11-13 min after the application of HM (Fig.3A-B). The second depolarization event was triggered with transient anoxia under both aCSF and HM incubation 15-20 min later. This caused propagating rSD under aCSF (Fig. 2A), but induced predominantly SiD (in total 35 out of 44 slices, 80%) in HM (Fig. 2B). In a few cases (n=3), SiD in HM was spontaneous, like SD1 (Suppl. Fig. 1A). Repolarization after SD1 in HM was substantially delayed or incomplete, and small amplitude, irregular field oscillations evolved in synchrony on the two DC channels over the repolarization phase (Fig. 2B; Suppl. Fig. 1B). These field oscillations invariably occurred in case SiD emerged subsequently, and appeared to predict SiD. Intriguingly, similar simultaneous field oscillations were observed in a malignant hemispheric stroke patient prior to the occurrence of terminal, simultaneous depolarization (Fig. 7A).

**Figure 6.**
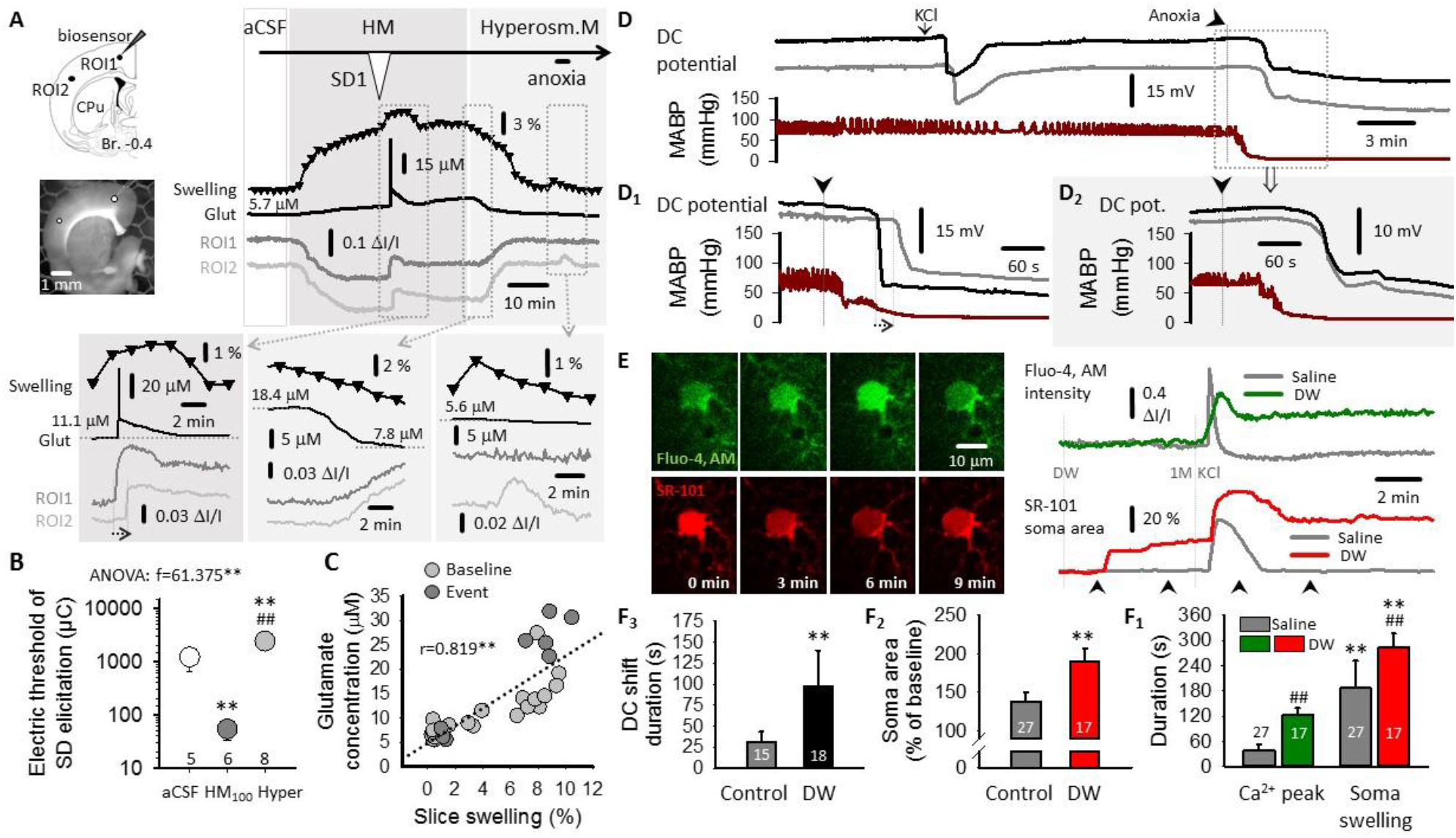
Reversal of slice swelling and SiD to SD with hyperosmotic medium (HRM), and the impact of water intoxication-induced cerebral edema on depolarization events and astrocyte calcium waves in anesthetized rodents. **A,** Changes of glutamate (Glut) accumulation and IOS signal (ROI1 & ROI2) after HRM treatment. **B,** The impact of HRM on the electrical threshold of SD elicitation, with respect to aCSF and HM_100_. **C,** Correlation between glutamate concentration and the degree of slice swelling. **D,** Representative DC potential traces (at two microelectrodes 3 mm apart), and mean arterial blood pressure – (MABP) after intraperitoneal (i.p.) distilled water (DW; D2) or physiological saline (D1) administration in anesthetized rats. Note the occurrence of SiD in the DW group. **E,** Two-photon imaging of intracellular calcium waves (Fluo-4, AM) in astrocytes, and astrocyte soma volume (SR-101) in the cerebral cortex of anesthetized mice. **F,** Quantitative analysis of the duration of the DC shift with SDs in anesthetized rats (F_1_), the degree of astrocytic soma swelling (F_2_), and the duration of the calcium wave and soma swelling associated with SDs (F_3_). Data in B, C, and F are expressed as mean±stdev; the number of analyzed depolarization events are shown in the charts. Statistical analysis relied on a one-way ANOVA (B), a one-tailed Pearson correlation analysis (C), a two-way ANOVA (F_1_) or an independent samples T test (F_2-3_); p<0.05* and p<0.01**. A Sidak post hoc test was applied whenever relevant; p<0.01** vs. aCSF, p<0.01## vs. HM_100_ (B); F_Ca/swell_=389.811**, F_DW_=216.143**, p<0.01** vs. Ca; p<0.01## vs. saline (F_1_).

**Figure 7.**
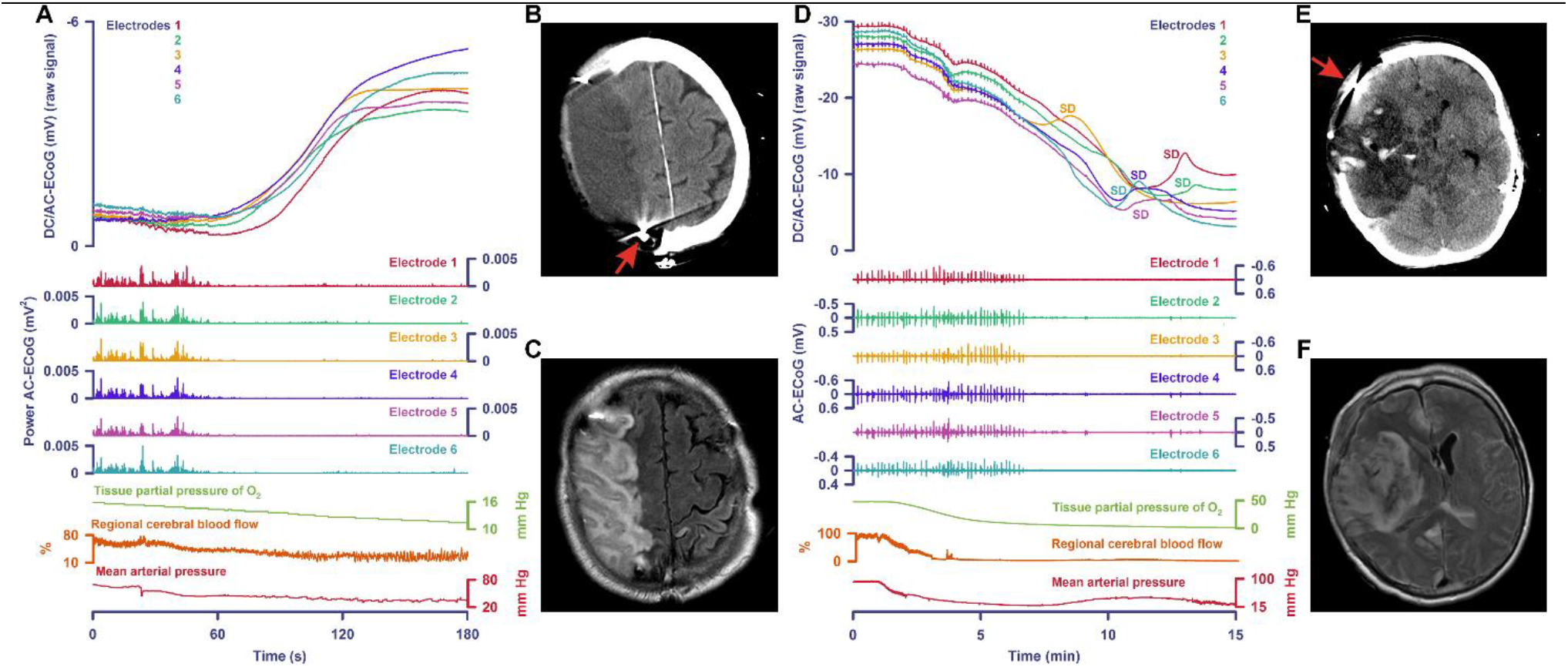
Terminal SiD and terminal SD in the human brain. Both patients were enrolled at Charité – Universitätsmedizin Berlin in a research protocol of invasive neuromonitoring approved by the local ethics committee, as published previously (Dreier *et al.*, 2018). **A-C** Terminal depolarization with very short depolarization arrival differences between different electrodes during dying in the wake of circulatory arrest. Decompressive hemicraniectomy was performed on day 1 because of malignant hemispheric stroke. A subdural electrode strip was implanted at the border zone between infarct and viable tissue. On day 4, neuroimaging showed several newly developed infarcts in other vascular territories and a Do Not Resuscitate-Comfort Care (DNRCC) order was activated followed by terminal extubation on day 5. **A**, Recordings of the direct current (DC)/alternate current (AC) electrocorticogram (ECoG) (frequency band: 0 - 45 Hz, upper 6 traces), the power of the AC band from 0.5 – 45 Hz at the 6 different electrodes (traces 7 – 12), brain tissue partial pressure of oxygen (p_ti_O_2_, intracortical Licox, Integra, Plainsboro, USA, trace 13), regional CBF (opto-electrode strip for laser-Doppler flowmetry, Perimed, Järfälla, Sweden, trace 14), and mean arterial pressure (radial artery catheter, trace 15). The changes during the final 180 seconds of the patient’s life are displayed. Parallel decreases of mean arterial pressure, rCBF and p_ti_O_2_ first result in non-spreading depression of spontaneous activity (traces 7-12). The terminal depolarization follows the complete non-spreading depression after 13 seconds. Note the almost simultaneous start of terminal depolarization at the 6 different electrodes in traces 1-6 suggesting SiD. Note, however, that in the exceptional condition that an SD propagates vertical to the electrode strip, “simultaneous depolarization” would also be perceived. **B**, Computed tomography (CT) reveals the infarct in the right middle cerebral artery territory. An electrode of the subdural electrode strip above the posterior part of the infarct is visible in this sectional plane (arrow). This is located almost exactly on the boundary between necrotic and living tissue. Typical streak artefacts are seen around the electrode (arrow). **C**, T2-weighted fluid-attenuated inversion recovery (FLAIR) magnetic resonance imaging (MRI) shows the infarct with higher accuracy. Electrodes are invisible in MRI scans. **D-F** Terminal SD during dying in the wake of circulatory arrest in a patient with subarachnoid and intracerebral hemorrhage. In addition, MRI on day 1 revealed large infarcts in several vascular territories. On day 2, a DNRCC order was activated and the patient was terminally extubated. **D**, Traces are similar to **A**, but traces 7-12 display simple spontaneous activity (0.5 – 45 Hz). The neuromonitored changes during the final 15 minutes of the patient’s life are displayed. Non-spreading depression of activity occurs during the falling phase of p_ti_O_2_. A very slow, homogeneous DC positivity started simultaneously with the fall in p_ti_O_2_. This is assumed to result from interferences of p_ti_O_2_ and pH with the electrode material (Dreier *et al.*, 2019). Superimposed on the DC positivity, terminal SD then started at electrode 3 and spread to the other electrodes. **E**, CT shows the condition after surgical removal of the large intracerebral hemorrhage. An electrode of the subdural electrode strip above living tissue of the frontal lobe is visible in this sectional plane (arrow). **F**, FLAIR image showing lesion and living tissue with higher accuracy.

The spatial resolution of the events was studied by replicating the same protocol with IOS imaging. Because IOS essentially reflects the swelling of the tissue, it was validated for depolarization events in a few slices by the synchronous recording of the DC potential (Suppl. Fig. 2A). The focus of SD1, which escaped visualization in the anesthetized rat (Fig. 1D_1_), was punctual (<1 % of total cortical area) in both aCSF and HM, and was localized to the upper layers of the dorsal-dorsolateral parietal cortex (Fig. 2C&E). SD1 propagated in the upper cortical layers (Fig. 2C), although the rate of propagation was significantly more rapid in HM than in aCSF (3.17±0.58 vs. 1.97±0.18 mm/min, HM vs. aCSF) (Fig. 2G). In aCSF, rSD evolved in response to anoxia as SD1, and spread at a rate of 2.48±0.31 mm/min (Fig. 2G). In contrast, anoxia in HM gave rise to SiD in 11 of 16 slices, implicating a sizeable area of the cortex (>50% of the total cortical area), which incorporated much of the tissue previously involved in SD1 propagation (Fig. 2D&F). In addition to the upper cortical layers, SiD progressively engaged the underlying deeper cortex, as well (Fig. 2D). Also, a propagating wave of SD took off at the ventral edge of SiD, and traveled towards the ventral tip of the cortex (Fig. 2D). Finally, in the remaining 5 of 16 slices, typically 3-4, spatially distinct, punctual SD foci emerged in temporal synchrony (cumulative focal area < 3 % of total cortical area) in response to anoxia, to give rise to propagating SD waves, which ultimately fused (Suppl. Fig. 2B). We identified these events as rSD with multifocal origin, and multifocal rSD was considered as a transition between classic rSD (i.e. single focus) and SiD.

Collectively, these data show that under osmotic stress, the tissue engaged in the propagation of SD1 later turned into a sizeable depolarization focus identified as SiD. The elevated IOS intensity sustained after SD1 (Fig. 2F_1_) and the concomitant failure of complete repolarization from SD1 (Suppl. Fig. 2) together predicted SiD occurrence, and were suggestive of tissue edema irreversible after SD1. After establishing a link between osmotic stress and SiD occurrence, we set out to explore the reciprocal interaction between tissue swelling and SD/SiD, because SD has been proposed as the principal mechanism of neuronal cytotoxic edema (Dreier et al., 2018; Kirov et al., 2020).

### Astrocyte swelling is implicated in SiD

To further explore the hypothesis that tissue swelling kindles SiD, we evaluated the degree of tissue swelling caused by the osmotic stress. We also examined the additional contribution of cytotoxic edema associated with SD/SiD to the overall tissue swelling. To this end, we first followed and measured changes in the surface area of brain slice preparations during the experimental protocol.

The increase of the brain slice area was most conspicuous with the second depolarization event, rSD in aCSF and SiD in HM (Fig. 3A_1-2_). Under aCSF, slice surface area was unchanging over time preceding SD1, then increased slightly with SD1 (with 1.09±0.68 pp.), and returned to baseline within minutes after SD1 (0.19±0.15 pp.) (Fig. 3A_3_&B). The maximum increase of slice area was greater with rSD than with SD1 (4.75±1.33 vs. 1.09±0.68 pp., rSD vs. SD1) (Fig. 3A_3_&B), but the swelling was again transient and reversible (Fig. 3A_3_). The transient swelling of the brain slice with SD1 and rSD as seen here is accepted to correspond to the cytotoxic edema previously associated with SD (Dreier et al., 2018). In HM, slice swelling commenced upon HM exposure, and the slice area gradually increased to reach a considerable expansion already before SD1 (4.59±1.14 and 4.15±1.11 vs. 0.07±0.09 pp., HM60 and HM_100_ vs. aCSF) (Fig. 3B). SD1 in HM occurred then spontaneously, and increased slice area with an additional, small extent (6.9±2.6 and 5.94±2 pp. HM60 and HM_100_). The slice area showed negligible variations afterwards, with no recovery to pre-SD1 level (Fig. 3A_4_&B). It is important that slice swelling preceded SD1 and SiD in HM, in contrast with aCSF, in which slice swelling was associated with SD1 and rSD.

Next, we set out to identify the cellular components of the profound macroscopic slice swelling. Since astrocytes rapidly swell in response to hypo-osmotic stress (Risher et al., 2009) we hypothesized the pivotal role of astroglial edema in our HM model. To confirm this concept, we examined astrocyte morphology in representative brain slices stained with the Golgi-Cox method, or in samples prepared for electron microscopy (Gull et al., 2015). Astrocytes appeared clearly swollen in both preparations after exposure to the hypo-osmotic medium, especially after the passage of SD1 (Fig. 3C). In the electron microscopic images, both the perinuclear plasma and astrocyte processes increased in size. The astrocyte plasma displayed decreased electron density indicative of cellular edema, particularly after SD1 had been superimposed to the osmotic stress (Fig. 3C_2_)

Consequently, we posited further that swollen astrocytes must substantially contribute to SiD, because astrocyte swelling increases neuronal excitability (Chebabo et al., 1995). We characterized the excitability of the nervous tissue by the amplitude of EFPs, the latency of SD/SiD onset with respect to anoxia onset, and the electric threshold of SD elicitation. EFP amplitude measured in the cerebral cortex was significantly greater in HM (171.0±11.8 and 184.3±7.4 vs. 93.1±9.3 μV, HM_60_ and HM_100_ vs. aCSF) (Fig. 3D). The latency of SiD occurrence to anoxia onset decreased markedly in HM (46.9±24.3 and 45.9±18.2 vs. 86.9±28.7 s; SiD in HM_60_ and HM_100_ vs. rSD in aCSF) (Fig. 3E). Likewise, the electric threshold of SD elicitation was drastically reduced in HM (50.1±15.7 and 52.5±18.4 vs. 1214.3±470.6 μC, HM_60_ and HM_100_ vs. aCSF) (Fig. 3F). In the latter set of experiments, the implication of astrocyte swelling was confirmed by repeating the SD threshold measurements with the addition of fluorocitrate to the aCSF, which decreased SD threshold similar to HM (51.0±22.9 and 52.5±18.4 μC, fluorocitrate and HM_100_) (Fig. 3F). We also explored whether impaired astrocyte metabolism caused by fluorocitrate is sufficient for SiD to occur. Fluorocitrate treatment replicated the HM phenotype: SD1 generated spontaneously in 4 of 5 slices, and SiD evolved in response to subsequent anoxia in 3 of 5 slices (Fig. 3G). Finally, both fluorocitrate and HM caused swelling of cortical astrocytes (Suppl. Fig. 3). Taken together, astrocyte swelling and dysfunction directly lead to bulk neuronal activation and created the conditions for SiD to occur.

### Astrocyte swelling is linked to SiD evolution to promote infarct maturation

Astrocyte swelling is pathogenic to the immediately adjacent tissue in cerebrovascular disease states (Kimelberg, 2005). Further, SD has been proposed to recruit viable ischemic penumbra tissue into the infarcted core (Hartings et al., 2017). Given this context, we hypothesized that astrocyte swelling and the related SiD increased the tissue volume at risk of injury. Moreover, we proposed that SiD must have represented the actual tissue infarction in progress, acknowledging that in other conditions, other types of SD may also cause neuronal death.

These suggestions were corroborated by our observation that the maximum depolarized tissue area engaged in SiD was invariably greater than the tissue area traversed by SD1 (76.8±11.4 vs. 60.9±10.7 %, SiD vs. SD1) (Fig. 4A). Furthermore, neurons, astrocytes, and the tissue at large were injured in the wake of SiD (Fig. 4B-D). In particular, electron microscopic images of the cortical area of SiD origin in a brain slice revealed widespread necrosis of neurons, occasionally seen in the close proximity of swollen astrocytes (Fig. 4B). Golgi-Cox staining identified astrocyte somata with extensively swollen contours and vestigial processes (Fig. 4D_1_), suggestive of advanced degeneration as compared to astrocyte morphology prior to SiD (Fig. 3C). The ultrastructural features of these astrocytes included prominent swelling of their soma, perivascular endfeet and mitochondria, and fragmented intracellular organelles (Fig. 4B&D_2_).

Macroscopic tissue damage assessed with TTC staining was obvious after SiD compared to SD1 in HM, or aCSF (Fig. 4C_1_). The quantitative evaluation of TTC staining showed that the number of TTC-positive cellular compartments (i.e. “particles”) was clearly reduced after SiD compared to SD1 in HM or aCSF (5.0±1.1 vs. 35.8±3.0 vs. 43.5±1.0 particles per 1000 μm^2^, SiD in HM vs. SD1 in HM vs. aCSF) (Fig. 4C_2_). Overall, the brain slices appeared to be severely damaged once they had suffered from SiD.

Complementary results were obtained from the brains of anesthetized rats. Twenty-five minutes after ischemia/anoxia related SiD occurrence, conventional Nissl-staining disclosed widespread neuronal necrosis in cortical brain slices (Fig. 4E). At the same time, the immunocytochemical co-localization of cleaved caspase-3 with NeuN or GFAP identified apoptotic astrocytes predominantly (Fig. 4F_1_). Cleaved caspase-3 was associated to the cell nucleus of astrocytes, and GFAP fibrils appeared to be fragmented in its close proximity (F_2_). Approximately 50 % of all GFAP-labeled astrocytes were cleaved caspase-3 positive in the parietal cortex, hippocampus and striatum, whereas only 4-12 % of NeuN-labeled neurons expressed cleaved caspase-3 (Fig. 4F_3_).

Our histological results together suggest that astrocytic edema and the linked SiD seriously jeopardize the survival of neurons and astrocytes alike, and impose significant damage to the nervous tissue.

### Inhibition of astrocyte swelling or volume regulated glutamate release blocks SiD

We have established above that a significant share of tissue swelling under osmotic stress was linked to the edema of astrocytes, which promoted SiD occurrence. In turn, SiD prognosticated acute damage to the nervous tissue. Next, we set out to understand the underlying mechanisms of SiD by attempting to prevent it by pharmacological means. Edema-related glutamate release or impaired glutamate clearance emerged as the most likely candidates to promote SiD, because astrocytes have been shown to swell and release glutamate in response to osmotic stress and spreading depolarization (Kimelberg et al. 1995, Basarsky et al., 1999, Yang et al. 2019). To test this notion, we first measured tissue glutamate concentration with enzymatic biosensors in anesthetized rats and live brain slice preparations, and pharmacologically modulated astrocyte swelling, the related glutamate release, glutamate receptor activation or glutamate clearance in brain slices.

As expected, SD1 after ischemia induction was associated with a sharp, transient rise of glutamate concentration in the anesthetized rat cortex (Fig. 5A). In contrast, the subsequent SiD in response to anoxia produced sustained glutamate accumulation (Fig. 5A). This pattern was reproducible in brain slices exposed to HM (Fig. 5B). Importantly, hypo-osmotic stress was associated with gradual glutamate accumulation to exceed 10 μM concentration by the time SD1 occurred. A sharp glutamate peak then took off coupled to SD1, but tissue glutamate concentration after recovery from SD1 remained constantly elevated above 15 μM, parallel with ongoing tissue swelling. SiD caused a second glutamate peak after which recovery was impaired, and glutamate concentration settled at a toxic level (>20 μM) against a background of marked edema (Fig. 5B).

Our pharmacological manipulations inhibited (i) astrocyte AQP4 channels and Na^+^/K^+^/Cl^-^ co-transporters (NKCCs) by TGN-020 + Bumetanide, or (ii) volume regulated anion channels (VRACs) by DCPIB, or (iii) antagonized NMDA and AMPA glutamate receptors by MK-801 + CNQX. All treatments were given in HM_60_. First, we evaluated the impact of each pharmacological treatment on slice swelling. TGN-020 + Bumetanide or DCPIB were both found very effective against tissue edema, as both treatments kept the tissue surface area significantly smaller than that under the HM_60_ condition (e.g. prior to SD1: 1.66±0.95 and 1.56±0.41 vs. 5.03±0.97 pp., TGN+Bum and DCPIB vs. HM_60_) (Fig. 5C). In contrast, MK-801+CNQX proved to be completely ineffective in this regard (e.g. prior to SD1: 4.68±0.04 vs. 5.03±0.97 pp., MK-801+ CNQX vs. HM_60_) (Fig. 5C). These data indicated that astrocytic AQP4 channels, NKCCs or VRACs must have played a pivotal role in tissue swelling in response to hypo-osmotic stress.

Next, we analyzed the typical features of depolarization events. Remarkably, all three treatments decreased the likelihood of SiD evolution very significantly. While SiD occurred in 28 of 33 slices (85 %) in HM_60_, SiD was seen in only 2 of 16 slices (13 %) in the TGN-020 + Bumetanide group, in 3 of 23 slices (13 %) in the DCPIB group, and 3 of 16 MK-801+CNQX-treated slices (19 %). Instead, the majority of the treated slices gave rise to rSD in response to anoxia. This was reflected in the size of the focal area of depolarization events. While the size of the SD1 focus was punctual and very small (<1 % of the cortical surface) in all groups (Fig. 5D), and the focus of SiD in the HM_60_ group engaged 55.5±7.2 % of the cortex as reported above, the rSD focus in the TGN+Bum and DCPIB groups was much more confined (1.8±0.7 and 3.9±0.7 vs. 55.5±7.2 %, rSD in TGN-Bum and DCPIB vs. SiD in HM_60_). The focal area of depolarization also decreased under glutamate receptor antagonism (9.4±4.0 vs. 55.5±7.2 %, rSD in MK-801+ CNQX vs. SiD in HM_60_), but this treatment was the least effective (Fig. 5D), and IOS imaging disclosed that rSD in the MK-801+ CNQX group was unique in that it was multifocal, as seen in a few HM slices before (Suppl. Fig. 2B). Taken together, the inhibition of astrocyte swelling by TGN-020 + Bumetanide (Igarashi et al., 2011; Kimelberg 2005) or DCPIB (Bourke et al., 1981) evidently blocked SiD evolution.

Finally, the impact of the pharmacological agents on tissue glutamate content was evaluated by measuring the area under the curve (AUC) during the glutamate peak with depolarization events (Fig. 5E-F). The glutamate load was invariably greater with rSD/SiD than with SD1, mainly due to the longer peak duration and impaired recovery after rSD/SiD. Both TGN-020 + Bumetanide and DCPIB application profoundly reduced tissue glutamate accumulation with SD1 (348.1±185.2 and 502.6±369.2 vs. 1416.9±757.6 μM × s, TGN020+Bum and DCPIB vs. HM_60_), and with rSD/SiD (1208.1±535.0 and 1324.6±983.2 vs. 4837.0±2081.7 μM × s, rSD in TGN020+Bum and DCPIB vs. SiD in HM_60_) (Fig. 5E). Glutamate receptor antagonism with MK-801+CNQX was again the least effective, achieving no significant reduction of glutamate load (e.g. 3196.1±1303.0 vs. 4837.0±2081.7 μM × s, rSD in MK-801+CNQX vs. SiD in HM_60_) (Fig. 5E). These results demonstrated that the reduction of astrocyte swelling by TGN-020 + Bumetanide or DCPIB, or the direct inhibition of volume-regulated glutamate release by DCPIB tempered the depolarization-related glutamate accumulation.

To further show that high glutamate levels lead to and are associated with SiD occurrence, we paralyzed glutamate uptake by the application of the excitatory amino acid transporter-2 (EAAT2) inhibitor TBOA in aCSF (Capuani et al., 2016) to some brain slices (Fig. 5F, n=4). Glutamate accumulated in these slices to 29.5±9.9 μM prior to the spontaneous occurrence of an SiD (3 of 4 slices), reached a peak concentration of 65.3±18.1 μM with SiD, and failed to recover afterwards, demonstrated by a glutamate level of 63.4±37.0 μM after SiD (Fig. 5F). These data further substantiated that glutamate accumulation (due to intensified release or impaired uptake) and must have mediated SiD in our osmotically challenged brain slices.

### The reversal of tissue swelling with a hyperosmotic medium restores physiological glutamate concentration and prevents SiD

Since the attenuation of astrocyte swelling and volume regulated glutamate release effectively decreased the susceptibility of the nervous tissue to SiD, we hypothesized that the complete reversal of cellular edema by hyperosmotic treatment could restore glutamate levels to the physiological range, and prevent SiD. To test this hypothesis, after 30 min incubation in HM and the spontaneous occurrence of SD1, HM was substituted with a hyperosmotic medium (HRM, 80 mM mannitol in aCSF), and the slice was challenged with anoxia as above. Slice edema and glutamate concentration elevation with SD1 replicated what was seen in our experiments presented above (Fig. 6A left and Fig. 5B). The perfusion of HRM over the slices completely reversed slice swelling as shown by the reduced slice area, as well as the recovery of the IOS signal. Parallel with the recovery from edema, HRM also restored physiological tissue glutamate concentration (Fig. 6A center). Further, anoxia did not trigger any depolarization event, and glutamate accumulation was not observed, either (Fig 7A right). Further, the electrical threshold of SD elicitation was markedly increased under HRM (2422.5±341.7 vs. 54.0±2.1 vs. 1220.0±6.2 μC, HRM vs. HM_100_ vs. aCSF) (Fig. 6B). The coincidence between slice swelling and tissue glutamate concentration was further supported by their good linear positive correlation (r=0.819) (Fig. 6C). We concluded that tissue glutamate content was tightly coupled to tissue edema, and that anti-edema treatment effectively counteracted tissue glutamate accumulation and SiD.

### Recapitulation of astrocyte swelling and SiD in the in vivo water intoxication model of cytotoxic edema

With the brain slice data in hand, we revisited our original assumption that cerebral edema is the precondition for SiD in anesthetized rodents (Fig. 1). To this end, we used the water intoxication model of cerebral edema, and recorded the cortical DC potential. Similar to the brain slices, a propagating event elicited experimentally with KCl was followed by a terminal SiD in response to anoxia in 6 of 7 rats. In the control condition (i.e. saline i.p.), the terminal event upon anoxia was a classic SD in 4 of 4 rats, as expected (Farkas et al., 2008). The KCl-induced SD superimposed on cerebral edema lasted considerably longer than a similar SD in control rats (94.4±48.3 vs. 30.4±11.1 s, water intoxication vs. saline) (Fig. 6F_1_), which stood in agreement with the brain slice recordings testifying that SiD in HM was preceded with an SD1 much longer than SD1 in aCSF (Suppl. Fig. 1C).

In order to underpin our brain slice observations that the swelling of astrocytes contributed significantly to cerebral edema (Fig. 3), we visualized astrocyte swelling (SR-101) and astrocyte Ca^2+^ dynamics (Fluo-4, AM) in anesthetized mice with two photon microscopy. We found that astrocyte somata became swollen after the initiation of water intoxication (in contrast with the level baseline in the control group), without detectable intracellular Ca^2+^ accumulation. Then, a KCl-triggered SD was accompanied with remarkable transient swelling of astrocytes, together with a matching, sharp rise of intracellular Ca^2+^ concentration (Fig. 6E). The SD-related astrocyte swelling was superimposed on the ongoing cerebral edema caused by water intoxication. Therefore a significantly greater degree of astrocyte swelling was reached compared to the control SD, which was reflected by the maximum surface area of selected astrocyte somata (160.47±12.29 vs. 134.85 %, water intoxication vs. control) (Fig. 6F_2_). The SD-related astrocyte swelling and intracellular Ca^2+^ accumulation lasted considerably longer in the water intoxicated group (soma swelling: 279.2±44.9 vs. 159.4±23.3 s, water intoxication vs. control; Ca^2+^ peak: 123.3±17.1 vs. 33.0±8.7 s, water intoxication vs. control) (Fig. 6F_3_). Finally, in contrast with full recovery seen in the control group, neither astrocyte soma volume, nor the intracellular concentration of Ca^2+^ recovered to pre-SD level in the water intoxicated group (Fig. 6E). We suggested that the duration of astrocyte swelling corresponded to the ability of the tissue to recover from SD. To conclude, we postulated that irreversible astrocyte swelling and dysfunction played a key role in the pathophysiology of SiD evolution.

## Discussion

The key findings of this study are summarized as follows. Cerebral edema is a sufficient condition for SD to occur spontaneously, which predisposes the tissue covered by the propagating SD for an upcoming simultaneous depolarization (SiD). The tissue volume engaged in the SiD then serves as an extensive focus of an SD event taking off from its perimeter. In particular, SiD is predicted by the swelling of astrocytes in edematous tissue. Astrocyte swelling under hypo-osmotic stress is implicated in the excessive extracellular accumulation of glutamate, coincident with SiD occurrence. Further, SiD is followed by the oncotic cell death of neurons and astrocytes, suggestive of infarct maturation associated with SiD. Finally, hyperosmotic intervention reduces the susceptibility of the nervous tissue to SD, and fully prevents the cascade of events leading to SiD.

This is the first study to characterize SiD comprehensively. An early review by Marshall (1959) did raise the possibility of SiD encompassing the entire cortex, but without providing any supportive experimental evidence. SiD-like events were later recorded in live brain slices incubated in hypo-osmotic medium and challenged with hypoxia (Kreisman et al., 2000) or in response to the bath application of supraphysiological concentration NMDA (Jarvis et al., 2001), but without detailed analysis of the phenomenon or realizing its significance. It is of high importance that an SiD-like event has been recently captured in the severely injured human brain (Dreier et al., 2018). In the ischemic penumbra of a malignant hemispheric stroke, terminal depolarization in the wake of circulatory arrest was seen to arrive with an unusual short delay from electrode to electrode on the subdural strip, when the strip was positioned at an already compromised area of the cortex (Fig. 7). This observation suggests that SiD is clinically relevant, and gives ample momentum to study the pathophysiological relevance of SiD in more detail.

SiD in our study occurred invariably superimposed on tissue edema, a condition to affect astrocytes first. The attention to astrocytes was also substantiated by our finding that fluorocitrate treatment caused astrocyte swelling and reproduced SiD evolution in brain slices. Unlike neurons, astrocytes are permeable to water because astrocytes are endowed with AQP-4 water channels at their processes, which conduct osmotically driven water (Andrew et al., 2007; Nagelhus et al., 2004). Further, astrocytes have been found to swell in response to ischemia (Wilson and Mongin, 2018) or SD (Risher et al., 2012), which is thought to represent water movement via AQP-4 along inward K^+^ currents. Alternatively, high extracellular K^+^ concentration – typical of SD – may initiate NKCC activation, which contributes to water accumulation within astrocytes (Kimelberg, 2005; Stokum et al., 2016; Wilson and Mongin, 2018). We found that HM-exposed astrocytes were markedly swollen in histological preparations, and that the blockade of AQP-4 channels and NKCCs prevented tissue edema and SiD occurrence under hypo-osmotic stress. These data collectively demonstrate that astrocyte swelling must be central to SiD evolution.

Astrocytic swelling has been implicated in volume regulated glutamate release relevant for ischemic stroke, through glutamate-permeable VRACs (Jayakumar and Norenberg, 2010; Wilson et al., 2019; Yang et al., 2019). At the same time, the significant glutamate uptake through astrocytic Na^+^- and ATP-dependent EAAT2 is impaired under metabolic stress, which sustains high extracellular glutamate concentration (Szatkowski et al., 1990; Rossi et al., 2000; Mahmoud et al., 2019). The inhibition of VRACs in our experiments prevented the accumulation of glutamate, reduced the focal area of depolarization and the likelihood of SiD occurrence. Conversely, EAAT2 blockade alone reproduced the SiD phenotype as seen under HM incubation. The antagonism of AMPA and NMDA receptors was partially effective against SiD, and resulted in multifocal SD evolution. These results suggest that surplus glutamate of astrocyte origin, in addition to neuronal release, must be implicated in the evolution of SiD. This seems plausible taken that glutamate appears to contribute to SD propagation (Pietrobon and Moskowitz, 2014), although several arguments cast doubt on the role of glutamate as the mediator driving the propagation of the SD wave front (Enger et al., 2015; Mei et al., 2020).

Previous observations suggest that even with only a very low residual blood flow, SD must last longer than 10 min before cell death occurs (Lückl et al., 2018). Our histological results suggest that this time span is significantly shortened when astrocytic function is already disturbed before the onset of neuronal depolarization. Thus, although SD is a primarily neuronal phenomenon (Peters et al., 2003; Chuquet et al., 2007), our findings substantiate the outstanding importance of astrocytes as protective guarantors against the devastating effects of SD (Largo et al., 1996; Dreier et al., 2015).

The clinical management of cerebral edema in acute brain injury currently aims at the reduction of intracranial pressure and the maintenance of cerebral perfusion pressure by sedation, hyperventilation, osmotherapy, hypothermia, and in the most severe cases decompressive craniectomy (Bardutzky and Schwab, 2007; Carney et al., 2017; Jha et al., 2019). However, for example, in severe subarachnoid hemorrhage intravenous high sodium fluids are administered increasingly more frequntly because hypoosmolarity is suggested to increase the risk and severity of delayed strokes (Schupper et al., 2020). Our results that preventive hyperosmotic intervention reduced the excitability of the nervous tissue and most importantly, averted SiD, provide pathophysiological insight into this empirical clinical strategy for the first time and emphasize the need to invent new ways of preventive osmotherapy in the treatment of acute brain injury.

## Funding

This work was supported by grants from the National Research, Development and Innovation Office of Hungary (No. PD128821 to ÁM, K134377 to EF, K120358 to FB, K135425 to IAK, FK132638 to AEF); the Ministry of Human Capacities of Hungary (ÚNKP-20-3-SZTE-110 to RF); the Economic Development and Innovation Operational Programme in Hungary co-financed by the European Union and the European Regional Development Fund (No. GINOP-2.3.2-15-2016-00048 to EF and GINOP-2.3.2-15-2016-0034 to IAK); the EU-funded Hungarian grant No. EFOP-3.6.1-16-2016-00008 to EF; the Deutsche Forschungsgemeinschaft (DFG DR 323/5-1 and DFG DR 323/10-1 to JPD), FP7 No. 602150 CENTER-TBI to JPD and Era-Net Neuron EBio2, with funds from BMBF (0101EW2004) to JPD; Inserm U1028 and CNRS UMR 5292 to AM and SM; grants from Fondation Les Gueules Cassées Sourire Quand Même (No. FGC 34-2018, FGC 49-2016 to AM and SM).

## Competing interests

The authors report no competing interests.

## Notes

### Competing Interest Statement

The authors have declared no competing interest.

## References

1. Andrew RD, Labron MW, Boehnke SE, Carnduff L, Kirov SA. Physiological evidence that pyramidal neurons lack functional water channels. Cereb Cortex. 2007 Apr;17(4):787–802.

2. Bardutzky J, Schwab S. Antiedema therapy in ischemic stroke. Stroke. 2007 Nov;38(11):3084–94.

3. Basarsky T A, Feighan D, and MacVicar BA. Glutamate Release through Volume-Activated Channels during Spreading Depression. J Neurosci. 1999 Aug 1; 19(15): 6439–6445.

4. Bere Z, Obrenovitch TP, Kozák G, Bari F, Farkas E. Imaging reveals the focal area of spreading depolarizations and a variety of hemodynamic responses in a rat microembolic stroke model. J Cereb Blood Flow Metab. 2014 Oct;34(10):1695–705.

5. Bourke RS, Waldman JB, Kimelberg HK, Barron KD, San Filippo BD, Popp AJ, Nelson LR. Adenosine-stimulated astroglial swelling in cat cerebral cortex in vivo with total inhibition by a non-diuretic acylaryloxyacid derivative. J Neurosurg. 1981 Sep;55(3):364–70.

6. Capuani C, Melone M, Tottene A, Bragina L, Crivellaro G, Santello M, Casari G, Conti F, Pietrobon D. Defective glutamate and K+ clearance by cortical astrocytes in familial hemiplegic migraine type 2. EMBO Mol Med. 2016 Aug 1;8(8):967–86.

7. Carney N, Totten AM, O’Reilly C, Ullman JS, Hawryluk GW, Bell MJ, Bratton SL, Chesnut R, Harris OA, Kissoon N, Rubiano AM, Shutter L, Tasker RC, Vavilala MS, Wilberger J, Wright DW, Ghajar J. Guidelines for the Management of Severe Traumatic Brain Injury, Fourth Edition. Neurosurgery. 2017 Jan 1;80(1):6–15.

8. Chebabo SR, Hester MA, Aitken PG, Somjen GG. Hypotonic exposure enhances synaptic transmission and triggers spreading depression in rat hippocampal tissue slices Brain Res. 1995 Oct 16;695(2):203–16.

9. Chuquet J, Hollender L, Nimchinsky EA. High-resolution in vivo imaging of the neurovascular unit during spreading depression. J Neurosci. 2007 Apr 11;27(15):4036–44.

10. Dreier JP. The role of spreading depression, spreading depolarization and spreading ischemia in neurological disease Nat Med. 2011 Apr;17(4):439–47.

11. Dreier JP, Fabricius M, Ayata C, Sakowitz OW, Shuttleworth CW, Dohmen C, Graf R, Vajkoczy P, Helbok R, Suzuki M, Schiefecker AJ, Major S, Winkler MK, Kang EJ, Milakara D, Oliveira-Ferreira AI, Reiffurth C, Revankar GS, Sugimoto K, Dengler NF, Hecht N, Foreman B, Feyen B, Kondziella D, Friberg CK, Piilgaard H, Rosenthal ES, Westover MB, Maslarova A, Santos E, Hertle D, Sánchez-Porras R, Jewell SL, Balança B, Platz J, Hinzman JM, Lückl J, Schoknecht K, Schöll M, Drenckhahn C, Feuerstein D, Eriksen N, Horst V, Bretz JS, Jahnke P, Scheel M, Bohner G, Rostrup E, Pakkenberg B, Heinemann U, Claassen J, Carlson AP, Kowoll CM, Lublinsky S, Chassidim Y, Shelef I, Friedman A, Brinker G, Reiner M, Kirov SA, Andrew RD, Farkas E, Güresir E, Vatter H, Chung LS, Brennan KC, Lieutaud T, Marinesco S, Maas AI, Sahuquillo J, Dahlem MA, Richter F, Herreras O, Boutelle MG, Okonkwo DO, Bullock MR, Witte OW, Martus P, van den Maagdenberg AM, Ferrari MD, Dijkhuizen RM, Shutter LA, Andaluz N, Schulte AP, MacVicar B, Watanabe T, Woitzik J, Lauritzen M, Strong AJ, Hartings JA. Recording, analysis, and interpretation of spreading depolarizations in neurointensive care: Review and recommendations of the COSBID research group J Cereb Blood Flow Metab. 2017 May;37(5):1595–1625.

12. Dreier JP, Lemale CL, Kola V, Friedman A, Schoknecht K. Spreading depolarization is not an epiphenomenon but the principal mechanism of the cytotoxic edema in various gray matter structures of the brain during stroke. Neuropharmacology. 2018 May 15;134(Pt B):189–207.

13. Dreier JP, Major S, Foreman B, Winkler MKL, Kang EJ, Milakara D, Lemale CL, DiNapoli V, Hinzman JM, Woitzik J, Andaluz N, Carlson A, Hartings JA. Terminal spreading depolarization and electrical silence in death of human cerebral cortex Ann Neurol. 2018 Feb;83(2):295–310.

14. Dreier JP, Major S, Lemale CL, Kola V, Reiffurth C, Schoknecht K, Hecht N, Hartings JA, Woitzik J. Correlates of Spreading Depolarization, Spreading Depression, and Negative Ultraslow Potential in Epidural Versus Subdural Electrocorticography. Front Neurosci. 2019 Apr 24;13:373.

15. Dreier JP, Reiffurth C. The stroke-migraine depolarization continuum. Neuron. 2015 May 20;86(4):902–922.

16. Enger R, Tang W, Vindedal GF, Jensen V, Johannes Helm P, Sprengel R, Looger LL, Nagelhus EA. Dynamics of Ionic Shifts in Cortical Spreading Depression. Cereb Cortex. 2015 Nov;25(11):4469–76.

17. Farkas E, Bari F, Obrenovitch TP. Multi-modal imaging of anoxic depolarization and hemodynamic changes induced by cardiac arrest in the rat cerebral cortex. Neuroimage. 2010 Jun;51(2):734–42.

18. Farkas E, Pratt R, Sengpiel F, Obrenovitch TP. Direct, live imaging of cortical spreading depression and anoxic depolarisation using a fluorescent, voltage-sensitive dye. J Cereb Blood Flow Metab. 2008 Feb;28(2):251–62.

19. Gull S, Ingrisch I, Tausch S, Witte OW, Schmidt S. Consistent and reproducible staining of glia by a modified Golgi-Cox method. J Neurosci Methods. 2015 Dec 30;256:141–50.

20. Hartings JA, Shuttleworth CW, Kirov SA, Ayata C, Hinzman JM, Foreman B, Andrew RD, Boutelle MG, Brennan KC, Carlson AP, Dahlem MA, Drenckhahn C, Dohmen C, Fabricius M, Farkas E, Feuerstein D, Graf R, Helbok R, Lauritzen M, Major S, Oliveira-Ferreira AI, Richter F, Rosenthal ES, Sakowitz OW, Sánchez-Porras R, Santos E, Schöll M, Strong AJ, Urbach A, Westover MB, Winkler MK, Witte OW, Woitzik J, Dreier JP. The continuum of spreading depolarizations in acute cortical lesion development: Examining Leão’s legacy. J Cereb Blood Flow Metab. 2017 May;37(5):1571–1594.

21. Hertelendy P, Menyhárt Á, Makra P, Süle Z, Kiss T, Tóth G, Ivánkovits-Kiss O, Bari F, Farkas E. Advancing age and ischemia elevate the electric threshold to elicit spreading depolarization in the cerebral cortex of young adult rats. J Cereb Blood Flow Metab. 2017 May;37(5):1763–1775.

22. Igarashi, H., Huber, V.J., Tsujita, M., and Nakada, T. (2011). Pretreatment with a novel aquaporin 4 inhibitor, TGN-020, significantly reduces ischemic cerebral edema. Neurol Sci 32, 113–116.

23. Jarvis CR, Anderson TR, Andrew RD. Anoxic depolarization mediates acute damage independent of glutamate in neocortical brain slices. Cereb Cortex. 2001 Mar;11(3):249–59.

24. Jayakumar AR, Norenberg MD. The Na-K-Cl Co-transporter in astrocyte swelling. Metab Brain Dis. 2010 Mar;25(1):31–8.

25. Jha RM, Kochanek PM, Simard JM. Pathophysiology and treatment of cerebral edema in traumatic brain injury. Neuropharmacology. 2019 Feb;145(Pt B):230–246.

26. Kimelberg HK, Kettenmann H. Swelling-induced changes in electrophysiological properties of cultured astrocytes and oligodendrocytes. I. Effects on membrane potentials, input impedance and cell-cell coupling Brain Res. 1990 Oct 8;529(1-2):255–61.

27. Kimelberg HK, Rutledge E, Goderie S, Charniga C. Astrocytic swelling due to hypotonic or high K+ medium causes inhibition of glutamate and aspartate uptake and increases their release. J Cereb Blood Flow Metab. 1995 May;15(3):409–16.

28. Kimelberg HK. Astrocytic swelling in cerebral ischemia as a possible cause of injury and target for therapy. Glia. 2005 Jun;50(4):389–97.

29. Kirov SA, Fomitcheva IV, Sword J. Rapid Neuronal Ultrastructure Disruption and Recovery during Spreading Depolarization-Induced Cytotoxic Edema. Cereb Cortex. 2020 Sep 3;30(10):5517–5531

30. Klass A, Sánchez-Porras R, Santos E. Systematic review of the pharmacological agents that have been tested against spreading depolarizations J Cereb Blood Flow Metab. 2018 Jul;38(7):1149–1179.

31. Kreisman NR, Soliman S, Gozal D. Regional differences in hypoxic depolarization and swelling in hippocampal slices. J Neurophysiol. 2000 Feb;83(2):1031–8.

32. Largo C, Cuevas P, Somjen GG, Martín del Río R, Herreras O. The effect of depressing glial function in rat brain in situ on ion homeostasis, synaptic transmission, and neuron survival. J Neurosci. 1996 Feb 1;16(3):1219–29.

33. Largo C, Ibarz JM, Herreras O. Effects of the gliotoxin fluorocitrate on spreading depression and glial membrane potential in rat brain in situ. J Neurophysiol. 1997 Jul;78(1):295–307.

34. Larrosa B., Pastor J., Lopez-Aguado L., et. al.: A role for glutamate and glia in the fast network oscillations preceding spreading depression. Neuroscience 2006; 141: pp. 1057–1068.

35. Lauritzen M, Dreier JP, Fabricius M, Hartings JA, Graf R, Strong AJ. Clinical relevance of cortical spreading depression in neurological disorders: migraine, malignant stroke, subarachnoid and intracranial hemorrhage, and traumatic brain injury J Cereb Blood Flow Metab. 2011 Jan;31(1):17–35.

36. Lückl J, Lemale CL, Kola V, Horst V, Khojasteh U, Oliveira-Ferreira AI, Major S, Winkler MKL, Kang EJ, Schoknecht K, Martus P, Hartings JA, Woitzik J, Dreier JP. The negative ultraslow potential, electrophysiological correlate of infarction in the human cortex. Brain. 2018 Jun 1;141(6):1734–1752.

37. Mahmoud S, Gharagozloo M, Simard C, Gris D. Astrocytes Maintain Glutamate Homeostasis in the CNS by Controlling the Balance between Glutamate Uptake and Release. Cells. 2019 Feb 20;8(2):184.

38. Marshall WH. Spreading cortical depression of Leao. Physiol Rev. 1959 Apr;39(2):239–79.

39. Matsuura T, Bures J. The minimum volume of depolarized neural tissue required for triggering cortical spreading depression in rat. Exp Brain Res. 1971;12(3):238–49.

40. Mei YY, Lee MH, Cheng TC, Hsiao IH, Wu DC, Zhou N. NMDA receptors sustain but do not initiate neuronal depolarization in spreading depolarization. Neurobiol Dis. 2020 Sep 2;145:105071.

41. Menyhárt Á, Farkas AE, Varga DP, Frank R, Tóth R, Bálint AR, Makra P, Dreier JP, Bari F, Krizbai IA, Farkas E. Large-conductance Ca2+-activated potassium channels are potently involved in the inverse neurovascular response to spreading depolarization. Neurobiol Dis. 2018 Nov;119:41–52.

42. Menyhárt Á, Zölei-Szénási D, Puskás T, Makra P, Bari F, Farkas E. Age or ischemia uncouples the blood flow response, tissue acidosis, and direct current potential signature of spreading depolarization in the rat brain. Am J Physiol Heart Circ Physiol. 2017 Aug 1;313(2):H328–H337.

43. Mestre H, Du T, Sweeney AM, Liu G, Samson AJ, Peng W, Mortensen KN, Stæger FF, Bork PAR, Bashford L, Toro ER, Tithof J, Kelley DH, Thomas JH, Hjorth PG, Martens EA, Mehta RI, Solis O, Blinder P, Kleinfeld D, Hirase H, Mori Y, Nedergaard M. Cerebrospinal fluid influx drives acute ischemic tissue swelling Science. 2020 Mar 13;367(6483):eaax7171.

44. Nagelhus EA, Mathiisen TM, Ottersen OP. Aquaporin-4 in the central nervous system: cellular and subcellular distribution and coexpression with KIR4.1. Neuroscience. 2004;129(4):905–13.

45. Obrenovitch TP, Chen S, Farkas E. Simultaneous, live imaging of cortical spreading depression and associated cerebral blood flow changes, by combining voltage-sensitive dye and laser speckle contrast methods. Neuroimage. 2009 Mar 1;45(1):68–74.

46. Peters O, Schipke CG, Hashimoto Y, Kettenmann H. Different mechanisms promote astrocyte Ca2+ waves and spreading depression in the mouse neocortex. J Neurosci. 2003 Oct 29;23(30):9888–96.

47. Pietrobon D, Moskowitz MA. Chaos and commotion in the wake of cortical spreading depression and spreading depolarizations. Nat Rev Neurosci. 2014 Jun;15(6):379–93.

48. Risher WC, Andrew RD, Kirov SA. Real-time passive volume responses of astrocytes to acute osmotic and ischemic stress in cortical slices and in vivo revealed by two-photon microscopy. Glia. 2009 Jan 15;57(2):207–21.

49. Risher WC, Ard D, Yuan J, Kirov SA. Recurrent spontaneous spreading depolarizations facilitate acute dendritic injury in the ischemic penumbra. J Neurosci. 2010 Jul 21;30(29):9859–68.

50. Risher WC, Croom D, Kirov SA. Persistent astroglial swelling accompanies rapid reversible dendritic injury during stroke-induced spreading depolarizations. Glia. 2012 Nov;60(11):1709–20.

51. Roper SN, Obenaus A, Dudek FE. Osmolality and nonsynaptic epileptiform bursts in rat CA1 and dentate gyrus. Ann Neurol. 1992 Jan;31(1):81–5.

52. Rossi DJ, Brady JD, Mohr C. Astrocyte metabolism and signaling during brain ischemia. Nat Neurosci. 2007 Nov;10(11):1377-86

53. Rossi DJ, Oshima T, Attwell D. Glutamate release in severe brain ischaemia is mainly by reversed uptake. Nature. 2000 Jan 20;403(6767):316–21.

54. Schupper AJ, Eagles ME, Neifert SN, Mocco J, Macdonald RL. Lessons from the CONSCIOUS-1 Study. J Clin Med. 2020 Sep 14;9(9):E2970.

55. Seidel JL, Faideau M, Aiba I, Pannasch U, Escartin C, Rouach N, Bonvento G, Shuttleworth CW. Ciliary neurotrophic factor (CNTF) activation of astrocytes decreases spreading depolarization susceptibility and increases potassium clearance. Glia. 2015 Jan;63(1):91–103.

56. Somjen GG. Mechanisms of spreading depression and hypoxic spreading depression-like depolarization Physiol Rev. 2001 Jul;81(3):1065–96.

57. Stokum JA, Gerzanich V, Simard JM. Molecular pathophysiology of cerebral edema. J Cereb Blood Flow Metab. 2016 Mar;36(3):513–38.

58. Swanson RA, Graham SH. Fluorocitrate and fluoroacetate effects on astrocyte metabolism in vitro. Brain Res. 1994 Nov 21;664(1-2):94–100.

59. Szatkowski M, Barbour B, Attwell D. Non-vesicular release of glutamate from glial cells by reversed electrogenic glutamate uptake. Nature. 1990 Nov 29;348(6300):443–6.

60. Tang YT, Mendez JM, Theriot JJ, Sawant PM, López-Valdés HE, Ju YS, Brennan KC. Minimum conditions for the induction of cortical spreading depression in brain slices. J Neurophysiol. 2014 Nov 15;112(10):2572–9.

61. Vasylieva N, Maucler C, Meiller A, Viscogliosi H, Lieutaud T, Barbier D, Marinesco S. Immobilization method to preserve enzyme specificity in biosensors: consequences for brain glutamate detection. Anal Chem. 2013 Feb 19;85(4):2507–15.

62. Verhaegen MJ, Todd MM, Warner DS, James B, Weeks JB. The role of electrode size on the incidence of spreading depression and on cortical cerebral blood flow as measured by H2 clearance. J Cereb Blood Flow Metab. 1992 Mar;12(2):230–7.

63. von Bornstädt D, Houben T, Seidel JL, Zheng Y, Dilekoz E, Qin T, Sandow N, Kura S, Eikermann-Haerter K, Endres M, Boas DA, Moskowitz MA, Lo EH, Dreier JP, Woitzik J, Sakadžić S, Ayata C. Supply-demand mismatch transients in susceptible peri-infarct hot zones explain the origins of spreading injury depolarizations. Neuron. 2015 Mar 4;85(5):1117–31.

64. Wilson CS, Bach MD, Ashkavand Z, Norman KR, Martino N, Adam AP, Mongin AA. Metabolic constraints of swelling-activated glutamate release in astrocytes and their implication for ischemic tissue damage. J Neurochem. 2019 Oct;151(2):255–272.

65. Wilson CS, Mongin AA. Cell Volume Control in Healthy Brain and Neuropathologies. Curr Top Membr. 2018;81:385–455.

66. Yamaguchi M, Wu S, Ehara K, Nagashima T, Tamaki N. Cerebral blood flow of rats with water-intoxicated brain edema. Acta Neurochir Suppl (Wien). 1994;60:190–2.

67. Yang J, Vitery MDC, Chen J, Osei-Owusu J, Chu J, Qiu Z. Glutamate-Releasing SWELL1 Channel in Astrocytes Modulates Synaptic Transmission and Promotes Brain Damage in Stroke. Neuron. 2019 May 22;102(4):813–827.e6.

