## Supplementary material for "Malignant astrocyte swelling and impaired glutamate clearance drive the expansion of injurious spreading depolarization foci"

**Supplementary Material to Menyhárt et al., 2020**

**Supplementary video 1.** A voltage sensitive dye fluorescence image sequence of the parietal cortical surface of an anesthetized rat. The image sequence was generated by background subtraction, contrasting and the application of a 3-frame moving average. Note the simultaneous depolarization (brightening of the signal) arising instantaneously in response to anoxia.

**Supplementary video 2.** A representative raw voltage sensitive dye fluorescence image sequence of the parietal cortical surface of an anesthetized rat. Note the macroscopic movement of the tissue, corresponding to cerebral edema, upon oxygen withdrawal.

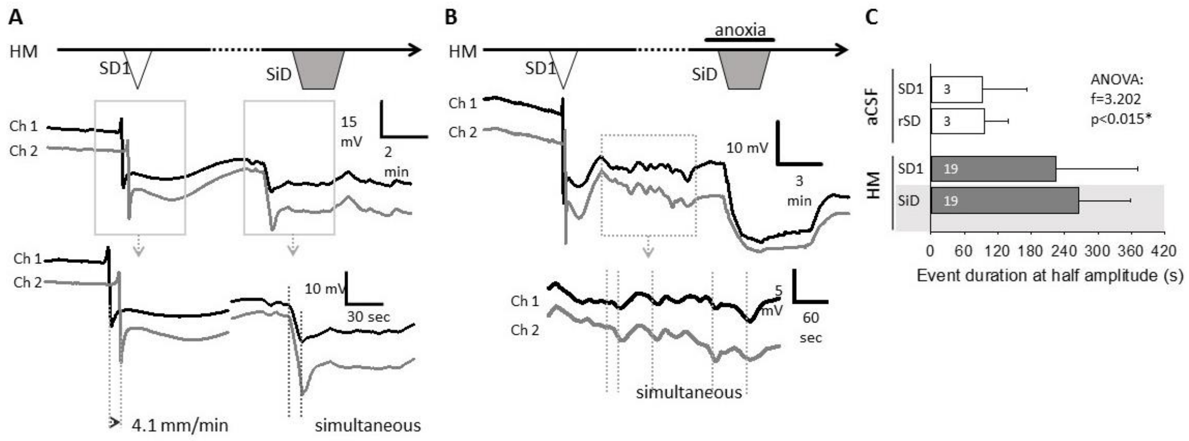


**Suppl. Fig. 1.** Direct current potential traces obtained with two microelectrodes (Ch 1 and Ch 2, about 1.3 mm apart) from representative live brain slice preparations exposed to hypo-osmotic medium (HM). **A,** Both the first spreading depolarization (SD1), and the subsequent simultaneous depolarization (SiD) occurred spontaneously, cancelling the need for the anoxic initiation of SiD. Note that the evolution of spontaneous SiD was similar to that of SiD triggered with transient anoxia in other slices. **B,** The repolarization phase of the first spreading depolarization (SD1) was incomplete, and small amplitude, irregular field oscillations evolved in synchrony on the two channels. This pattern of the DC signal was typical in case simultaneous depolarization (SiD) ensued. **C,** The duration of depolarization events taken at half amplitude of the negative DC potential shift. Both the first spreading depolarization (SD1) and recurrent SD (rSD)/simultaneous depolarization (SiD) lasted significantly longer in hypo-osmotic medium (HM) than in artificial cerebrospinal fluid (aSCF). Data are given as mean±stdev, the number of events analyzed is shown in each bar. A one-way ANOVA paradigm was used for statistical evaluation of the data ($p<0.05$*$p<0.01$**).

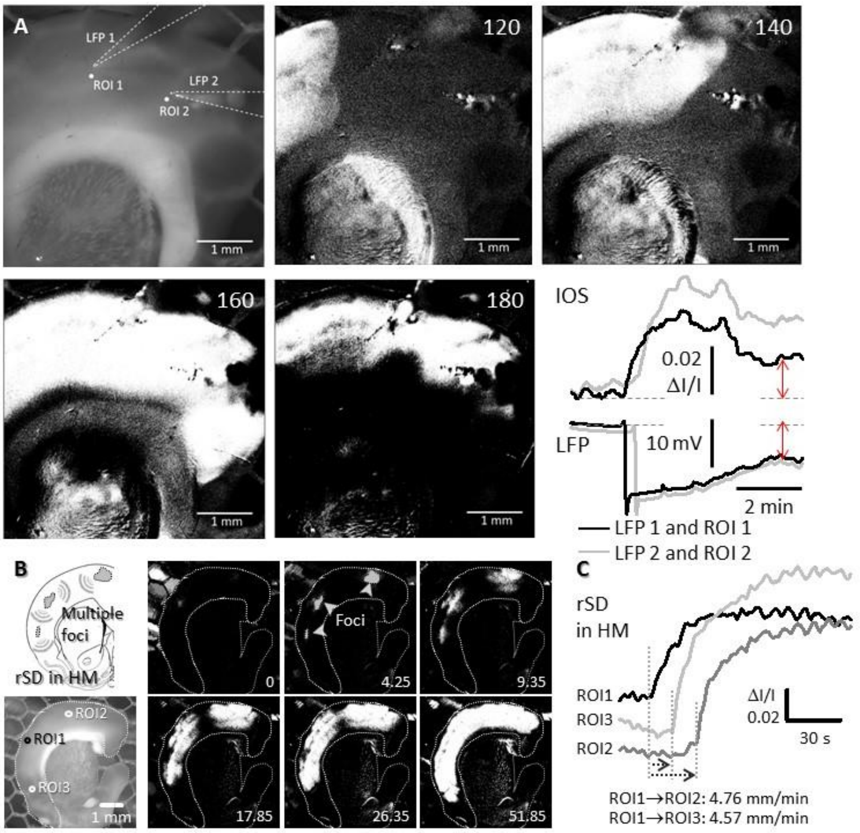


**Suppl. Fig. 2.** Intrinsic optical signal (IOS) signal imaging in live brain slice preparations. **A,** The correspondence between the intrinsic optical signal (IOS) intensity changes and the direct current potential recordings in a representative live brain slice preparation exposed to hypo-osmotic medium (HM). The evolution of a spreading depolarization event (SD1) that occurred spontaneously in HM is shown. The two regions of interest (ROI1-2) to be used for the extraction of the optical signal intensity changes were positioned adjacent to the two local field potential glass capillary microelectrodes (LFP1-2). The corresponding traces are given in the lower right hand corner. The temporal resolution of the image sequences is given in the upper right corner in seconds; time is displayed with respect to SD1 onset. **B,** Representative images of a recurrent spreading depolarization (rSD) triggered with transient anoxia in a live brain slice preparation exposed to hypo-osmotic medium ($HM_{100}$). Note that the event propagated from three foci, attesting a multi-focal origin. Intrinsic optical signal intensity changes at three regions of interest (ROI1-3) placed between the foci demonstrated the propagation of the rSD. Images were obtained by template matching (movement artefact correction), background subtraction, smoothing and contrast enhancement. The temporal resolution of the image sequences is given in the lower right corner in seconds.

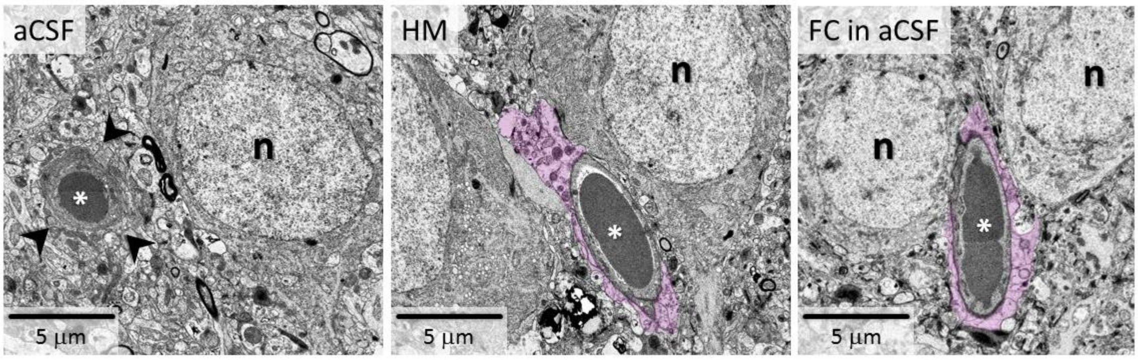


**Suppl. Fig. 3.** Representative electron microscopic images of astrocyte endfeet embracing cerebrocortical microvessels. The samples were taken from live brain slice preparations and show pyramidal cells (n) near the microvessels (*). While the perivascular environment appeared electron dense and intact in normal artificial cerebrospinal fluid (aCSF) (black arrowheads), the perivascular astrocyte endfeet (highlighted with purple) retained water and were swollen in hypo-osmotic medium (HM) and after fluorocitrate (FC) treatment equally.
